# SARS-CoV-2 nsp15 endoribonuclease antagonizes dsRNA-induced antiviral signaling

**DOI:** 10.1101/2023.11.15.566945

**Authors:** Clayton J. Otter, Nicole Bracci, Nicholas A. Parenti, Chengjin Ye, Li Hui Tan, Abhishek Asthana, Jessica J. Pfannenstiel, Nathaniel Jackson, Anthony R. Fehr, Robert H. Silverman, Noam A. Cohen, Luis Martinez-Sobrido, Susan R. Weiss

## Abstract

Severe acute respiratory syndrome coronavirus (SARS-CoV)-2 has caused millions of deaths since emerging in 2019. Innate immune antagonism by lethal CoVs such as SARS-CoV-2 is crucial for optimal replication and pathogenesis. The conserved nonstructural protein 15 (nsp15) endoribonuclease (EndoU) limits activation of double-stranded (ds)RNA-induced pathways, including interferon (IFN) signaling, protein kinase R (PKR), and oligoadenylate synthetase/ribonuclease L (OAS/RNase L) during diverse CoV infections including murine coronavirus and Middle East respiratory syndrome (MERS)-CoV. To determine how nsp15 functions during SARS-CoV-2 infection, we constructed a mutant recombinant SARS-CoV-2 (nsp15^mut^) expressing a catalytically inactive nsp15. Infection with SARS-CoV-2 nsp15 ^mut^ led to increased activation of the IFN signaling and PKR pathways in lung-derived epithelial cell lines and primary nasal epithelial air-liquid interface (ALI) cultures as well as significant attenuation of replication in ALI cultures compared to wild-type (WT) virus. This replication defect was rescued when IFN signaling was inhibited with the Janus activated kinase (JAK) inhibitor ruxolitinib. Finally, to assess nsp15 function in the context of minimal (MERS-CoV) or moderate (SARS-CoV-2) innate immune induction, we compared infections with SARS-CoV-2 nsp15^mut^ and previously described MERS-CoV nsp15 mutants. Inactivation of nsp15 had a more dramatic impact on MERS-CoV replication than SARS-CoV-2 in both Calu3 cells and nasal ALI cultures suggesting that SARS-CoV-2 can better tolerate innate immune responses. Taken together, SARS-CoV-2 nsp15 is a potent inhibitor of dsRNA-induced innate immune response and its antagonism of IFN signaling is necessary for optimal viral replication in primary nasal ALI culture.

**SIGNIFICANCE:** Severe acute respiratory syndrome coronavirus (SARS-CoV)-2 causes a spectrum of respiratory disease ranging from asymptomatic infections to severe pneumonia and death. Innate immune responses during SARS-CoV-2 infection have been associated with clinical disease severity, with robust early interferon responses in the nasal epithelium reported to be protective. Thus, elucidating mechanisms through which SARS-CoV-2 induces and antagonizes host innate immune responses is crucial to understanding viral pathogenesis. CoVs encode various innate immune antagonists, including the conserved nonstructural protein 15 (nsp15) which contains an endoribonuclease (EndoU) domain. We demonstrate that SARS-CoV-2 EndoU is a crucial interferon antagonist, by providing further evidence for the role of the conserved CoV nsp15 in antagonizing innate immune activation, thereby optimizing CoV replication.

## INTRODUCTION

There are currently seven known human coronaviruses (HCoVs), three of which emerged in the past 20 years to cause severe disease [severe acute respiratory syndrome (SARS)-CoV, Middle East respiratory syndrome (MERS)-CoV, and SARS-CoV-2]. SARS-CoV-2 is the causative agent of coronavirus disease 2019 (COVID-19) and the ongoing pandemic that has claimed millions of lives worldwide [1].

CoV genomes are positive-sense, single-stranded (ss)RNAs of approximately 30 kilobases in length. The 5’ proximal two-thirds of the CoV genome is composed of two open reading frames (ORFs), ORF1a and ORF1b, that encode 16 non-structural proteins (nsps), while the 3’ proximal third encodes the structural and accessory proteins. Following viral entry, ORF1a and ORF1b are translated from genomic RNA and processed into nsps that form replication-transcription complexes (RTCs) localized to double-membrane vesicles formed from rearranged endoplasmic reticulum. Nsps also function as RNA-processing enzymes, proteases, and protein modifiers [2]. Here, we focused on nsp15, which contains an endoribonuclease domain (EndoU) that is localized to the RTC during infection. We previously reported that, during infection of bone marrow-derived macrophages (BMMs), murine coronavirus nsp15 EndoU cleaves genomic RNA with a preference for U↓A and C↓A sequences [3]. Alternatively, it has been proposed that nsp15 EndoU cleaves polyU sequences on the 5′-end of negative-sense, antigenome RNA [4]. Both studies concluded that EndoU reduces viral double-stranded (ds)RNA, resulting in limited activation of dsRNA-induced antiviral responses, including interferon (IFN) signaling, the oligoadenylate synthetase/ribonuclease L (OAS/RNase L) pathway, and the protein kinase R (PKR) pathway [2, 3, 5].

DsRNA is a byproduct of viral replication that can be detected by host cell sensors. Melanoma differentiation-associated protein 5 (MDA5) detects dsRNA and triggers the production of type I and type III IFNs [6–9]. Once secreted, IFN binds to its receptor on both infected and uninfected cells, which results in Janus kinase (JAK) activation and phosphorylation of signal transducer and activator of transcription (STAT)1 and STAT2. STAT phosphorylation and subsequent nuclear translocation promotes transcription from promoters that regulate a coordinated antiviral response involving hundreds of interferon-stimulated genes (ISGs) [10–12]. Upon sensing of dsRNA, OASs (isoforms 1-3) produce 2’-5’-oligoadenylates (2-5A), which activate RNase L. RNase L cleaves both viral and cellular RNAs, thereby suppressing viral replication and protein synthesis as well as inducing inflammation and apoptosis [13, 14]. PKR autophosphorylates upon sensing of dsRNA and then phosphorylates eukaryotic initiation factor 2 (eIF2)a, leading to activation of the integrated stress response and inhibition of protein synthesis [15]. Both PKR and OASs are ISGs and, due to the interconnected nature of these pathways, dsRNA is a key regulator of many antiviral host cell responses.

Previous studies have characterized the endoribonuclease activity of nsp15 expressed by the murine coronavirus, mouse hepatitis virus (MHV) [3, 5, 16, 17]. Mutation of either catalytic histidine (H262A or H277A) within the EndoU domain of MHV nsp15 promotes increased induction of type I IFN, activation of the OAS/RNase L and PKR pathways, and significant attenuation of replication in both bone-marrow derived macrophages (BMMs) and in mice [16, 17]. These findings suggest that inactivation of EndoU results in increased sensing of dsRNA by host sensors which was concluded to occur due to either an increase in total dsRNA or a redistribution of dsRNA away from RTCs [3, 16]. Similar finding were reported with mutant viruses expressing catalytically inactive nsp15 of HCoV-229E, avian infectious bronchitis virus (IBV), porcine epidemic diarrhea virus (PEDV), and MERS-CoV [5, 16–20]. We have previously characterized multiple recombinant MERS-CoV mutant viruses with catalytically inactivated nsp15, deletion of accessory protein NS4a (a dsRNA-binding protein), inactivation of accessory protein NS4b (which contains a phosphodiesterase that specifically antagonizes RNase L activation and also function as an IFN signaling inhibitor via inhibition of the NF-κB pathway) or double mutants of both nsp15/NS4a or nsp15/NS4b [19, 21–24]. We found that although the MERS-CoV nsp15 mutant virus had a minor replication defect in respiratory epithelial cell lines, the double mutants were significantly more attenuated and elicited increased expression of IFN and ISG mRNAs [19]. These data suggest that nsp15 is crucial to CoV evasion of host detection due to its ability to antagonize dsRNA-induced innate immune responses.

Published reports on SARS-CoV-2 nsp15 have utilized either *in vitro* biochemical assays, ectopic overexpression, or bioinformatic approaches to characterize the structural and RNA-binding/cleavage capacity of SARS-CoV-2 nsp15 [25–34]. However, these studies were carried out in the absence of other viral proteins and moreover did not address the role of nsp15 in SARS-CoV-2 infection. Therefore, to understand the activity of nsp15 EndoU in the context of authentic infection, we generated and characterized a recombinant SARS-CoV-2 expressing a catalytically inactive nsp15 (SARS-CoV-2 nsp15^mut^). We assessed the impact of nsp15 on viral replication and host responses in lung-derived epithelial cell lines as well as primary nasal epithelial air-liquid interface (ALI) cultures, which recapitulate many features of the *in vivo* upper airway, including its heterogeneous cellular population (primarily ciliated epithelial cells and mucus-producing goblet cells) and mucociliary functions [35, 36]. These nasal ALI cultures mimic the initial site of infection, where innate immune responses are crucial for early control of viral infections [37, 38]. Our data indicate that SARS-CoV-2 nsp15 contributes to innate immune evasion by dampening IFN induction, ISG expression, and PKR pathway activation in all cell types examined, and that inactivation of nsp15 results in attenuation of replication in primary cell ALI cultures.

## RESULTS

### Generation of recombinant SARS-CoV-2 nsp15^mut^ expressing an inactive endoribonuclease

To determine the role of nsp15 in antagonizing dsRNA-induced innate immune responses during SARS-CoV-2 infection, we generated a recombinant SARS-CoV-2 expressing a catalytically inactive nsp15 (denoted SARS-CoV-2 nsp15^mut^) using a bacterial artificial chromosome (BAC)-based reverse genetic system [39]. SARS-CoV-2 nsp15 contains two catalytic histidines at amino acid residue positions 234 and 249, conserved among coronaviruses, as illustrated for a subset of betacoronaviruses in **Figure 1A** [40]. We generated a catalytically inactive nsp15 via a histidine (H) to alanine (A) substitution at amino acid position 234 (nsp15^H234A^). To validate that the H234A substitution was sufficient to abrogate endoribonuclease activity, nsp15 wild type (WT) and the H234A mutant proteins were expressed in *E. coli* and purified to homogeneity (**Figure S1**). Endoribonuclease activity of both nsp15 proteins was tested using a fluorescence-based assay in which a synthetic ssRNA substrate containing a Förster resonance energy transfer (FRET) pair was incubated with each recombinant nsp15. Endoribonuclease activity results in cleavage of ssRNA substrate and fluorescence detection (**Figures 1B and 1C**). While nuclease activity was observed for WT nsp15, the activity of nsp15^H234A^ was comparable to that of buffer alone. These results suggest that the H234A substitution in place of a catalytic histidine was sufficient to abolish the endoribonuclease activity of SARS-CoV-2 nsp15.

**Figure 1.**
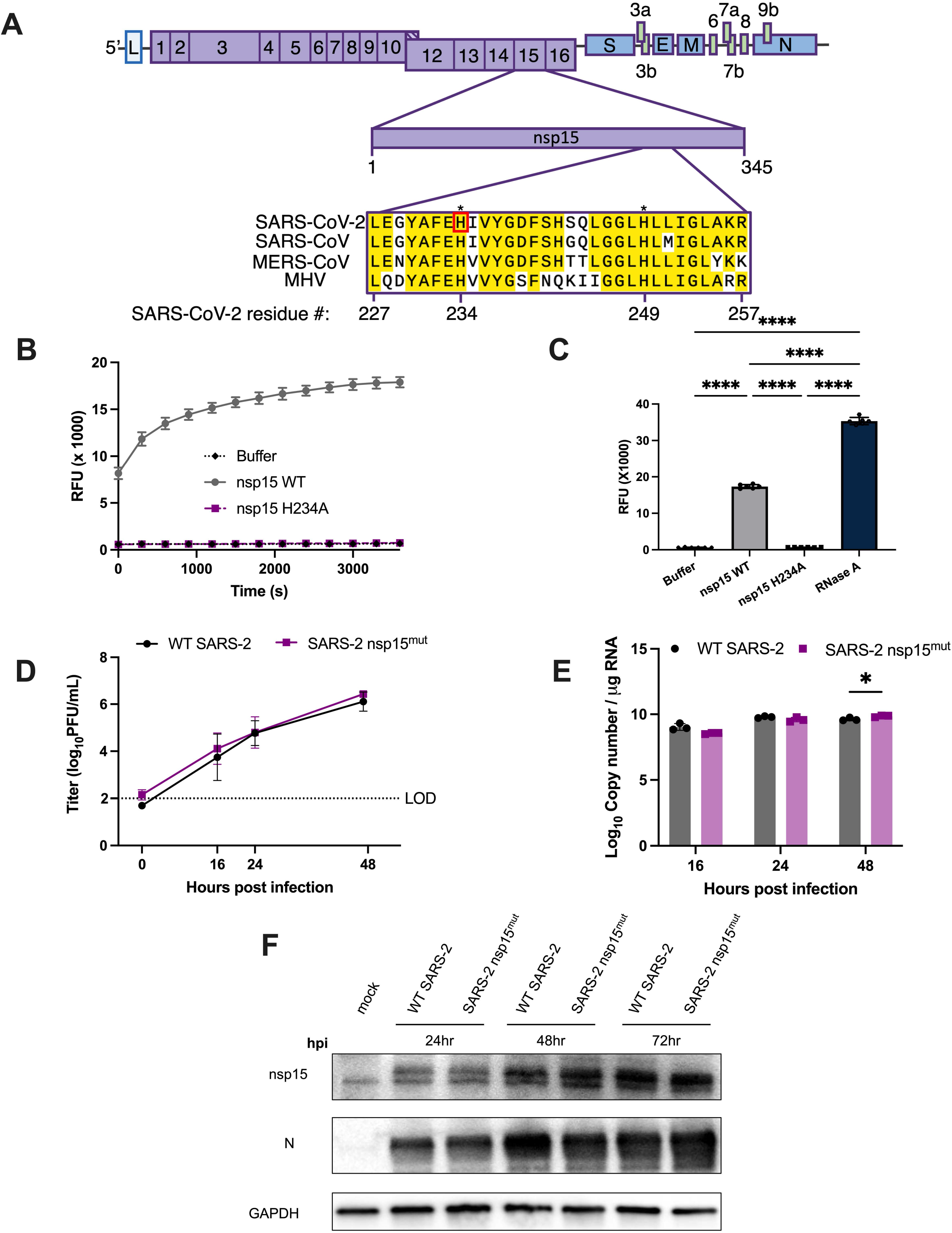
Construction of recombinant SARS-CoV-2 with inactive nsp15^mut^ endoribonuclease. (A) Diagram of the SARS-CoV-2 genome. Full-length nsp15 of SARS-CoV-2 is shown in the center. Sequence alignment among beta-coronaviruses, from top to bottom: SARS-CoV, SARS-CoV-2 USA-WA1/2020, MERS-CoV EMC/2012, and MHV strain A59. Conserved residues are shown in yellow with the catalytic histidine residues of SARS-CoV-2, H234 and H249, designated with asterisks. Nsp15 mutation site for SARS-CoV-2 nsp15^mut^ generated in this study, H234, shown in red. (B) SARS-CoV-2 WT, nsp15^mut^, and buffer alone catalytic activity measured by dequenching of 6-FAM upon ssRNA cleavage measured in relative fluorescence units (RFU) recorded at 5-minute intervals over a 60-minute period. (C) Total quantified ssRNA cleavage measured for the WT nsp15, nsp15^mut^, buffer alone, and RNase A at the end of the 1 hour. (D-F) Vero E6 cells were infected with either WT or nsp15^mut^ SARS-CoV-2 (MOI 0.1 in D and MOI 1 in E and F). (D) Supernatants were collected at indicated times post-infection and titered via plaque assay. Limit of detection (LOD) at 100 PFU/mL is indicated. Data shown is the average of 2 independent experiments. (E) Intracellular RNA was collected at the indicated time points post-infection and copies of the RdRp per μg RNA was measured by RT-qPCR targeting nsp12 using a standard curve. (F) Cells were lysed at indicated time points using protein lysis buffer. Samples were separated via SDS-PAGE and transferred to a PVDF membrane for immune detection with antibodies against SARS-CoV-2 nsp15, SARS-CoV-2 N, and GAPDH.

To determine if the H234A substitution was associated with an inherent viral growth defect, we compared the kinetics of replication of SARS-CoV-2 nsp15^mut^ with that of WT SARS-CoV-2 in VeroE6 cells, an IFN-deficient cell line [41, 42]. Viral growth curves quantifying infectious virus in the supernatant of infected cells at various time points post-infection revealed no significant difference between WT and nsp15^mut^ SARS-CoV-2 (**Figure 1D**). Corroborating this finding, no significant differences in viral genome copy number (quantified via RT-qPCR from intracellular RNA) were detected (**Figure 1E**). Additionally, western blots probed for nsp15 and nucleocapsid (N) protein revealed no differences in viral protein expression between WT and nsp15^mut^ SARS-CoV-2 (**Figure 1F**). Thus, the H234A catalytic mutation in SARS-CoV-2 nsp15^mut^ does not directly impact SARS-CoV-2 replication in IFN-incompetent VeroE6 cells.

### Infection with SARS-CoV-2 nsp15^mut^ promotes increased IFN signaling and PKR pathway activation compared to WT SARS-CoV-2 in respiratory epithelial cell lines

To understand the role that SARS-CoV-2 nsp15 plays in evading dsRNA-induced innate immunity, we utilized two lung-derived epithelial cell lines, A549 cells transduced to stably express the SARS-CoV-2 receptor, human angiotensin converting-enzyme 2 (ACE2) (A549-ACE2), and Calu3 cells. Both of these cell lines have fully intact IFN signaling, PKR pathway, and OAS/RNase L pathway responses. We and others have previously reported that WT SARS-CoV-2 induces all of these pathways during infection of both A549-ACE2 and Calu3 cells [9, 43–45].

Infection of either A549-ACE2 (**Figure 2A**) or Calu3 (**Figure 3A**) cells with SARS-CoV-2 nsp15^mut^ revealed no significant difference in replication kinetics compared to WT SARS-CoV-2 (**Figures 2A and 3A**). However, a slight but significant decrease in intracellular viral genome copies was detected at 48 and 72 hours post infection (hpi) during SARS-CoV-2 nsp15^mut^ infection compared to WT in Calu3 cells **(Figures 2B and 3B**).

**Figure 2.**
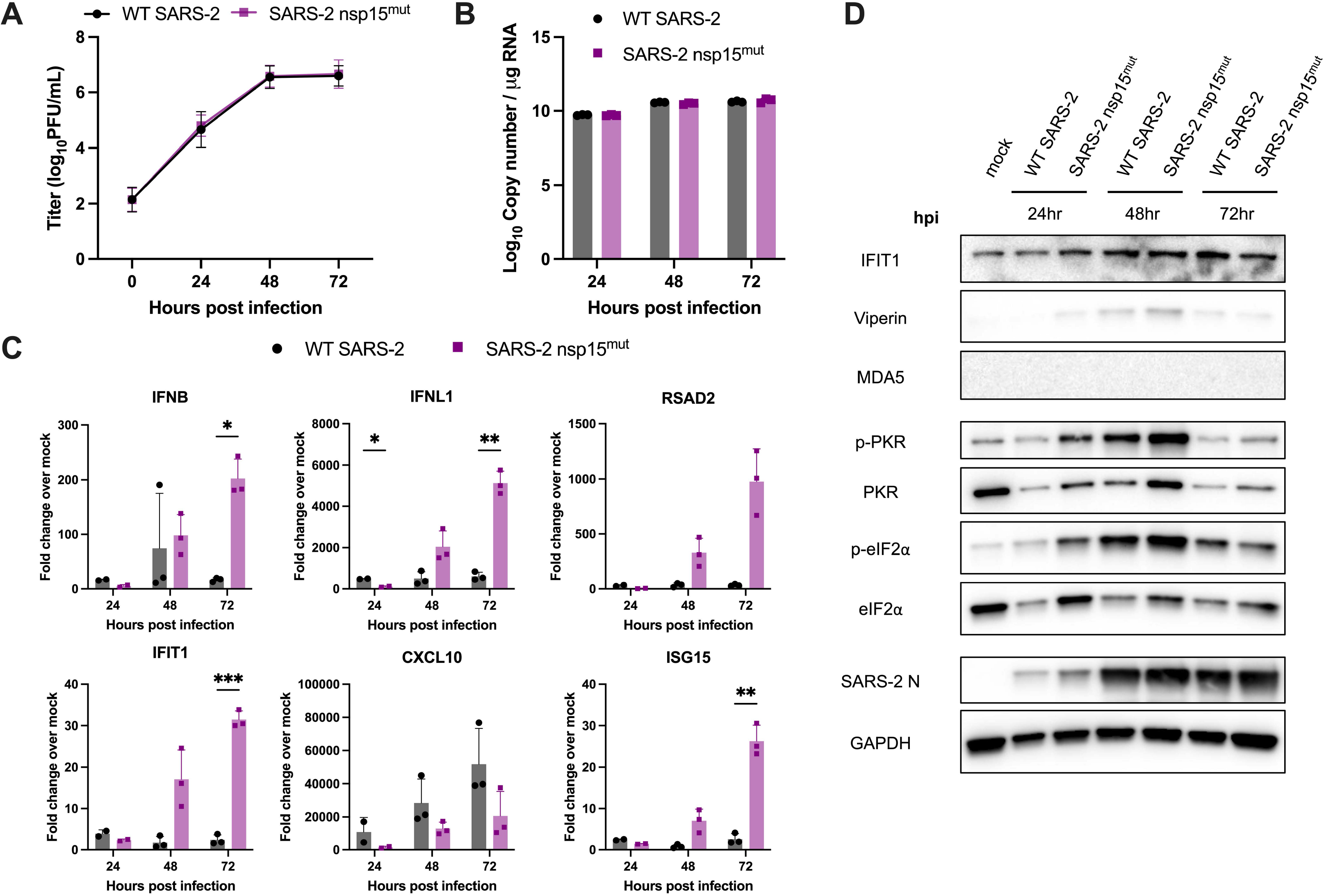
SARS-2-nsp15^mut^ induces increased IFN mRNA and ISG expression as well as PKR activation in A549-ACE cells compared to WT SARS-CoV-2. A549-ACE2 cells were infected with either WT or nsp15^mut^ SARS-CoV-2 at MOI 0.1 (A) or MOI 1 (B-D). (A) At indicated time points, supernatants were collected and infectious virus was measured by plaque assay. Data shown is the average of three independent experiments. (B-D) At 24, 48, and 72hpi either intracellular RNA was extracted and analyzed by RT-qPCR or whole cell lysates were collected. (B) Genome copy number was calculated using a standard curve and primers targeting SARS-CoV-2 RdRp. (C) Relative mRNA expression of *IFNLB, IFNL1*, and four representative ISGs: *RSAD2*, IFIT1 *CXCL10*, and *ISG15* were measured. Fold change is expressed as ΔΔCt. (D) Whole cell lysates were separated via SDS-PAGE and analyzed by western blot analysis with antibodies against indicated proteins. Data in B-D are from one representative experiment of three total experiments.

**Figure 3.**
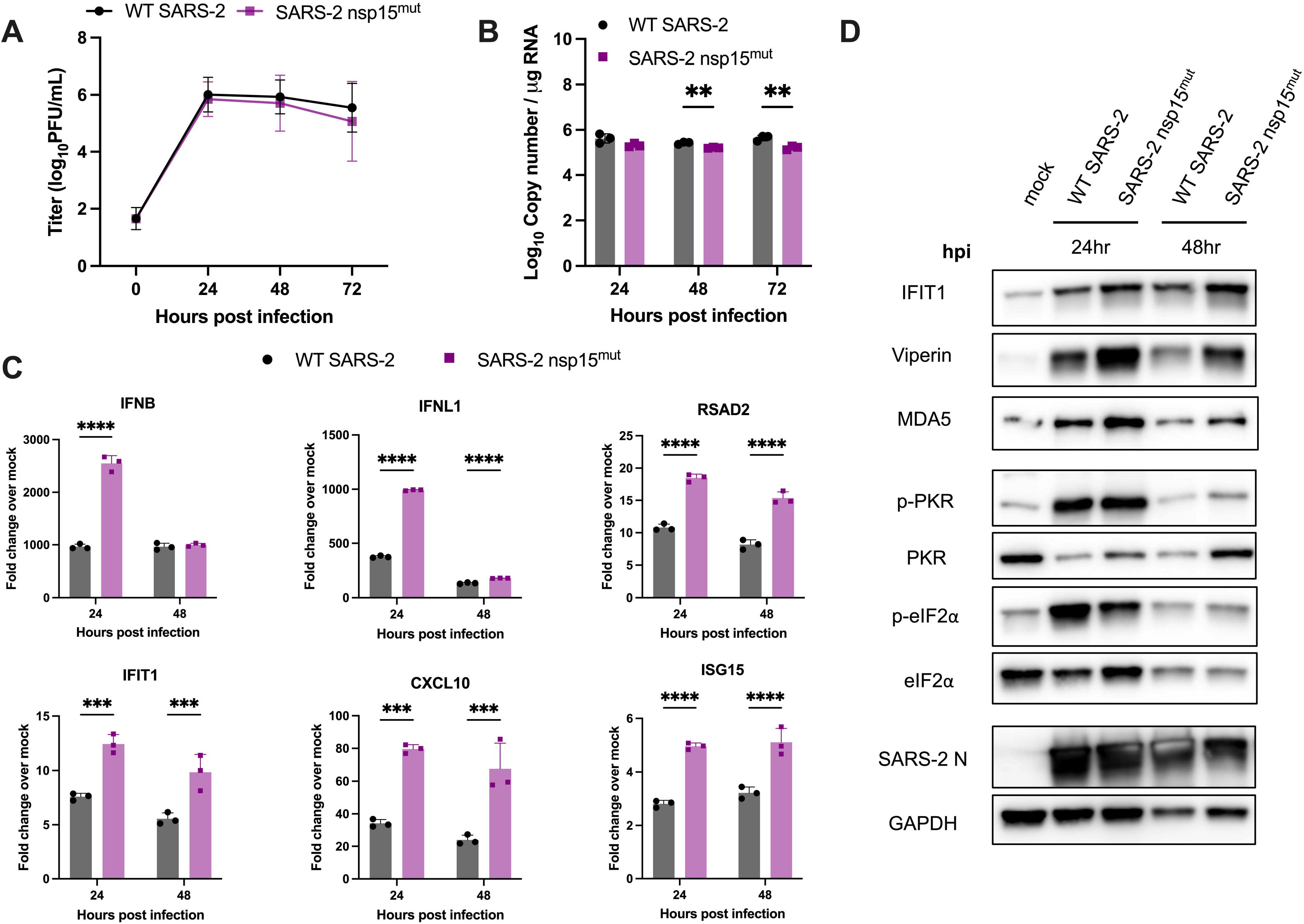
SARS-2-nsp15^mut^ induces increased IFN mRNA and ISG expression in Calu3 cells compared to WT SARS-CoV-2. Calu3 cells were infected with either WT or nsp15^mut^ SARS-CoV-2 (A) MOI 0.1 and (B-D) MOI 1). (A) Supernatants were collected at 0, 24, 48, and 72hpi and viral titers were measured by plaque assay. Data shown is the average of three independent experiments. (B-C) At the indicated time points post-infection, intracellular RNA was extracted and analyzed by RT-qPCR. (B) Genomes copy numbers were quantified using primers against SARS-CoV-2 nsp12 and a standard curve. (C) Relative mRNA expression of *IFNB, IFNL1*, and four representative ISGs: *RSAD2*, *IFIT1*, *CXCL10*, and *ISG15.* Fold change is expressed as ΔΔCt. D) Whole cell lysates were resolved using SDS-PAGE and analyzed via western blot analysis using antibodies against indicated proteins Data in B-D are from one representative experiment of three total experiments.

Expression of type I (*IFNB*) and type III IFN (*IFNL1*) mRNAs, as well as mRNA expression of four representative ISGs (radical S-adenosyl methionine domain containing 2 (RSAD2), IFN-induced protein with tetratricopeptide repeats 1 (*IFIT1*), C-X-C motif chemokine ligand 10 (*CXCL10)*, and *ISG15)* were quantified by RT-qPCR to detect activation of the IFN signaling pathway **(Figures 2C and 3C**). Despite the lack of viral attenuation, we observed increased mRNA induction of both type I and type III IFNs upon infection with SARS-CoV-2 nsp15^mut^ compared to WT in both cell lines. mRNA expression of representative ISGs was also upregulated in A549-ACE2 and Calu3 cells with SARS-CoV-2 nsp15^mut^.

To confirm our observations of increased IFN and ISG mRNA induction, western blots were performed to assess protein expression of IFIT1, viperin, MDA5, and PKR in lysates collected from cells either mock-infected or infected with WT or nsp15^mut^ SARS-CoV-2. While levels of IFIT1 and viperin were slightly increased during infection with nsp15^mut^ relative to WT early (24 hpi) during infection in A549-ACE2, MDA5 was not detected (**Figure 2D**). Similar analysis of infected Calu3 cells revealed a clear increase in IFIT1, viperin, MDA5, and PKR expression during SARS-CoV-2 nsp15^mut^ infection compared to WT at both 24 and 48 hpi (**Figure 3D**).

Activation of the PKR pathway was also evaluated via western blot to assess phosphorylation of PKR and its downstream substrate eIF2α. Both p-PKR and p-eIF2α levels were increased during SARS-CoV-2 nsp15^mut^ infection relative to WT in A549-ACE2 cells (**Figure 2D**) indicating both an earlier (24 hpi) and more robust activation of the PKR pathway during SARS-CoV-2 nsp15^mut^ infection. Interestingly, similar analysis in Calu3-infected cells revealed clear phosphorylation of PKR and eIF2α early (24 hpi) during infection, however phosphorylation levels of PKR and eIF2α were reduced at 48 hpi compared to 24 hpi in both WT and nsp15^mut^ infected cells. Though increased PKR pathway activation (above WT levels) was detected in A549-ACE2 cells, we hypothesize that robust activation of the PKR pathway by WT SARS-CoV-2 in Calu3 cells renders any additional PKR activation by SARS-CoV-2 nsp15^mut^ difficult to detect. It is important to note that in both cell lines, comparable levels of SARS-CoV-2 N were observed following infection with either nsp15^mut^ or WT SARS-CoV-2, consistent with the lack of replication defect **(Figures 2D and 3D**). Finally, we compared OAS/RNase L activation in A549-ACE2 and Calu3 cells infected with WT and nsp15^mut^ SARS-CoV-2 by assessing ribosomal RNA (rRNA) degradation on a bioanalyzer. No increase in RNase L activity was observed during SARS-CoV-2 nsp15^mut^ infection relative to activation observed by WT SARS-CoV-2 in either cell type (**Figures S2 and S3**).

### SARS-CoV-2 nsp15^mut^ is attenuated for replication and induces increased IFN signaling and PKR pathway activation in primary nasal epithelial ALI cultures

We sought to investigate the role of nsp15 in a primary nasal epithelial cell culture system which models the initial site of viral replication and the primary barrier to respiratory virus infections. Thus, nasal epithelial cells derived from four to six donors were pooled prior to growth and differentiation at an air-liquid interface (ALI) to recreate the cell types and functions present in the nasal airway. Infections comparing WT and nsp15^mut^ SARS-CoV-2 in nasal ALI cultures were conducted at 33°C (approximate nasal airway temperature) to optimize replication of SARS-CoV-2 and model the *in vivo* nasal temperature dynamics [46–48].

To investigate the kinetics of viral replication, nasal ALI cultures were infected at the apical surface with WT or nsp15^mut^ SARS-CoV-2. Apical surface liquid (ASL) was collected at 48-hour intervals from 0 to 192 hpi and quantified via plaque assay. Average viral titers are shown in **Figure 4A**. WT SARS-CoV-2 reached peak titers at 144 hpi and then plateaued, as we have previously reported [46]. In contrast to growth curves in both A549-ACE2 and Calu3 cells, SARS-CoV-2 nsp15^mut^ replication was significantly attenuated compared to WT by approximately 10-fold PFU/mL at both 144 and 192 hpi. In addition, a significant reduction in intracellular genome copy number was detected at both 96 and 192 hpi during SARS-CoV-2 nsp15^mut^ infection compared to WT as quantified by RT-qPCR (**Figure 4B**).

**Figure 4.**
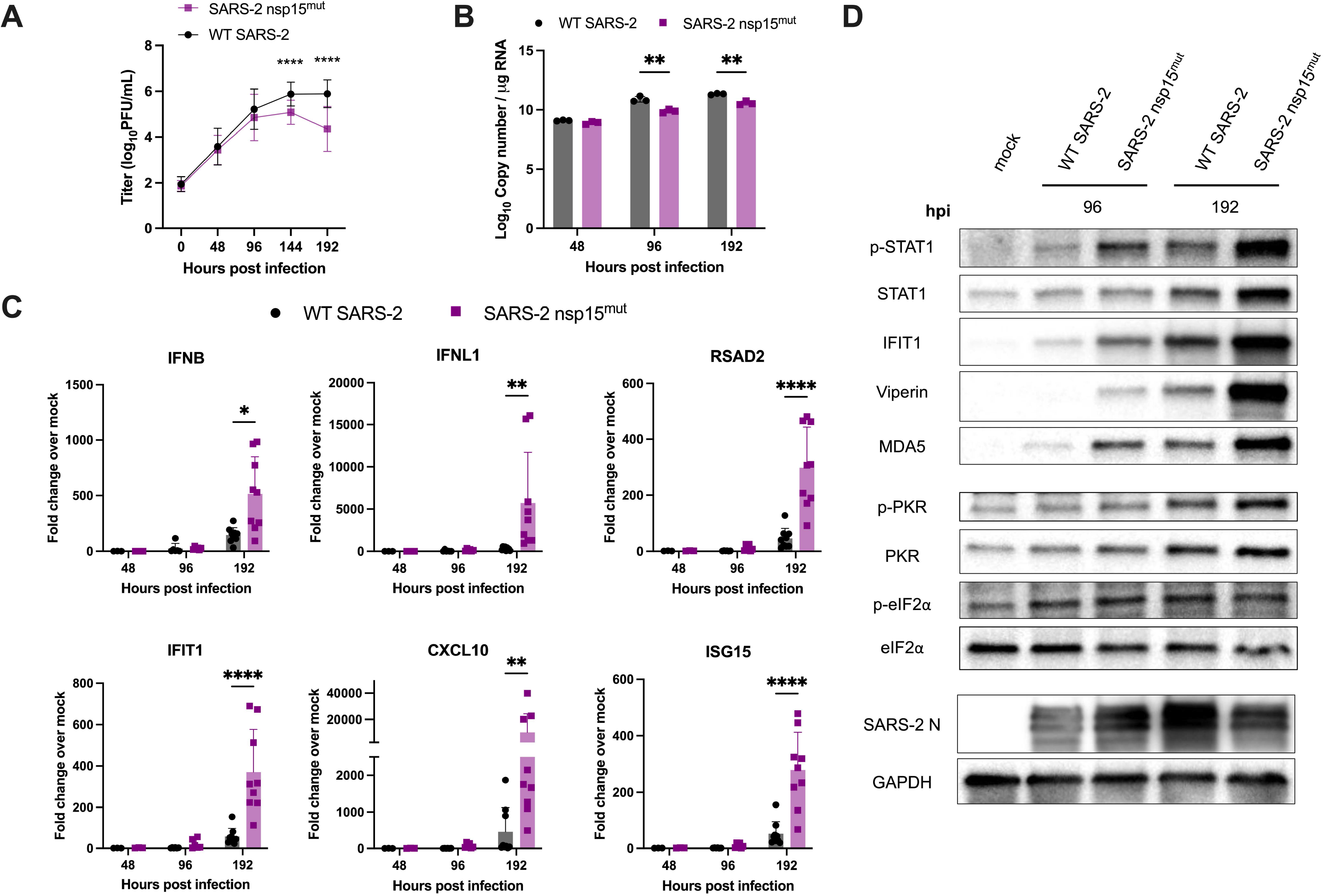
SARS-2-nsp15^mut^ is attenuated for replication compared to WT SARS-CoV-2 and induces increased IFN signaling and PKR pathway activation in primary nasal epithelial ALI cultures. Primary nasal ALI cultures were infected at an MOI of 1 with either WT or nsp15^mut^ SARS-CoV-2. (A) Apical surface liquid was collected every 48 hours and infectious virus was titered by plaque assay. Growth curves shown are the average titers from 6 independent experiments, each performed in triplicate with a different set of 4-6 pooled nasal cell donors. (B-C) Intracellular RNA was extracted and analyzed by RT-qPCR at 48, 96 and 192 hpi. B) Genomes copies were measured using primers against nsp12 and compared against a standard curve. (C) Relative mRNA expression of *IFNB* and *IFNL1* as well as four representative ISGs: *RSAD2, IFIT1 CXCL10*, and *ISG15*. Fold Change is expressed as ΔΔCt. (D) Western blot analysis of whole cell lysates collected at 96 and 192 hpi was performed using antibodies against indicated proteins. Data in B-D are from one representative experiment of 3 total experiments.

Next, we compared dsRNA-induced innate immune responses in nasal ALI cultures infected with either WT or nsp15^mut^ SARS-CoV-2. We observed delayed IFN and ISG responses during both WT and nsp15 mutant virus infections. At 48 and 96 hpi, there was minimal induction of IFN and ISG mRNA in cultures infected with either virus (**Figure 4C**). However, both IFN and the representative ISG mRNAs were significantly (∼10-fold) upregulated at 192 hpi during SARS-CoV-2 nsp15^mut^ infection compared to WT. The kinetics of IFN and ISG mRNA induction in nasal ALI cultures were consistent with the timing of the replication defect observed for SARS-CoV-2 nsp15^mut^ (**Figure 4A**).

Given the growth defect and increased expression of IFN and ISG mRNA during infection with SARS-CoV-2 nsp15^mut^ compared to WT in nasal ALI cultures, we examined STAT1 phosphorylation (p-STAT1) and ISG induction by western blot. At 96 hpi, we observed an increase in p-STAT1, as well as increased levels of ISG protein expression (IFIT1, Viperin, MDA5, and PKR) in samples infected with nsp15^mut^ compared to WT SARS-CoV-2 (**Figure 4D**). This increase in p-STAT1 and upregulation of ISGs during SARS-CoV-2 nsp15^mut^ infection was even more robust at 192 hpi.

Protein lysates from infected nasal ALI cultures were further evaluated to assess PKR pathway activation (**Figure 4D**). A clear increase in p-PKR as well as total PKR was observed during infection with both viruses at 192 hpi. Despite a clear increase in p-PKR, we have not detected a consistent increase in p-eIF2α above mock levels in nasal ALI cultures from 5 independent sets of pooled donors infected with either WT SARS-CoV-2 or the nsp15 mutant at any time point assayed. Western blotting for SARS-CoV-2 N levels showed decreased expression at 192 hpi during nsp15^mut^ infection compared to WT (**Figure 4D**), consistent with nsp15^mut^ attenuation in primary nasal cells.

### Attenuation of SARS-CoV-2 nsp15^mut^ in primary nasal epithelial cells is IFN-mediated

To determine the extent to which the attenuation of SARS-CoV-2 nsp15^mut^ virus is due to IFN signaling, we infected nasal ALI cultures in the presence of the small molecule JAK1/2 inhibitor, ruxolitinib (RUX). RUX treatment prevents phosphorylation of STATs by JAK1/2 and subsequent nuclear translocation; thus RUX-treated nasal cells can produce and release IFN, but cells are unable to respond to IFN and induce ISG transcription [49]. Nasal ALI cultures pre-treated with RUX were infected with either WT or nsp15^mut^ SARS-CoV-2. ASL was collected every 48 hours and infectious virus quantified via plaque assay. As expected, in control DMSO-treated cultures, replication of SARS-CoV-2 nsp15^mut^ was attenuated compared to WT at late times post infection (144 and 192 hpi) (**Figure 5A**). While RUX treatment led to a significant increase in replication of WT SARS-CoV-2 (ten-fold at 192 hpi), RUX treatment of SARS-CoV-2 nsp15^mut^ infected cultures resulted in a larger increase in viral titers (100-fold). However, no significant difference in replication was observed between the RUX treated cultures infected with either WT or nsp15^mut^ SARS-Cov-2 at 192 hpi. **Figure 5B** highlights the viral titers from **Figure 5A** for each virus and treatment condition at 192 hpi plotted as a bar graph, illustrating complete rescue of the nsp15^mut^ growth defect to WT SARS-CoV-2 levels with RUX treatment. As might be expected, RUX treatment significantly impacts viral titers at the late time points post infection, which coincides with the timing of IFN signaling induction during SARS-CoV-2 infection of nasal cells (**Figures 4C and 4D**).

**Figure 5.**
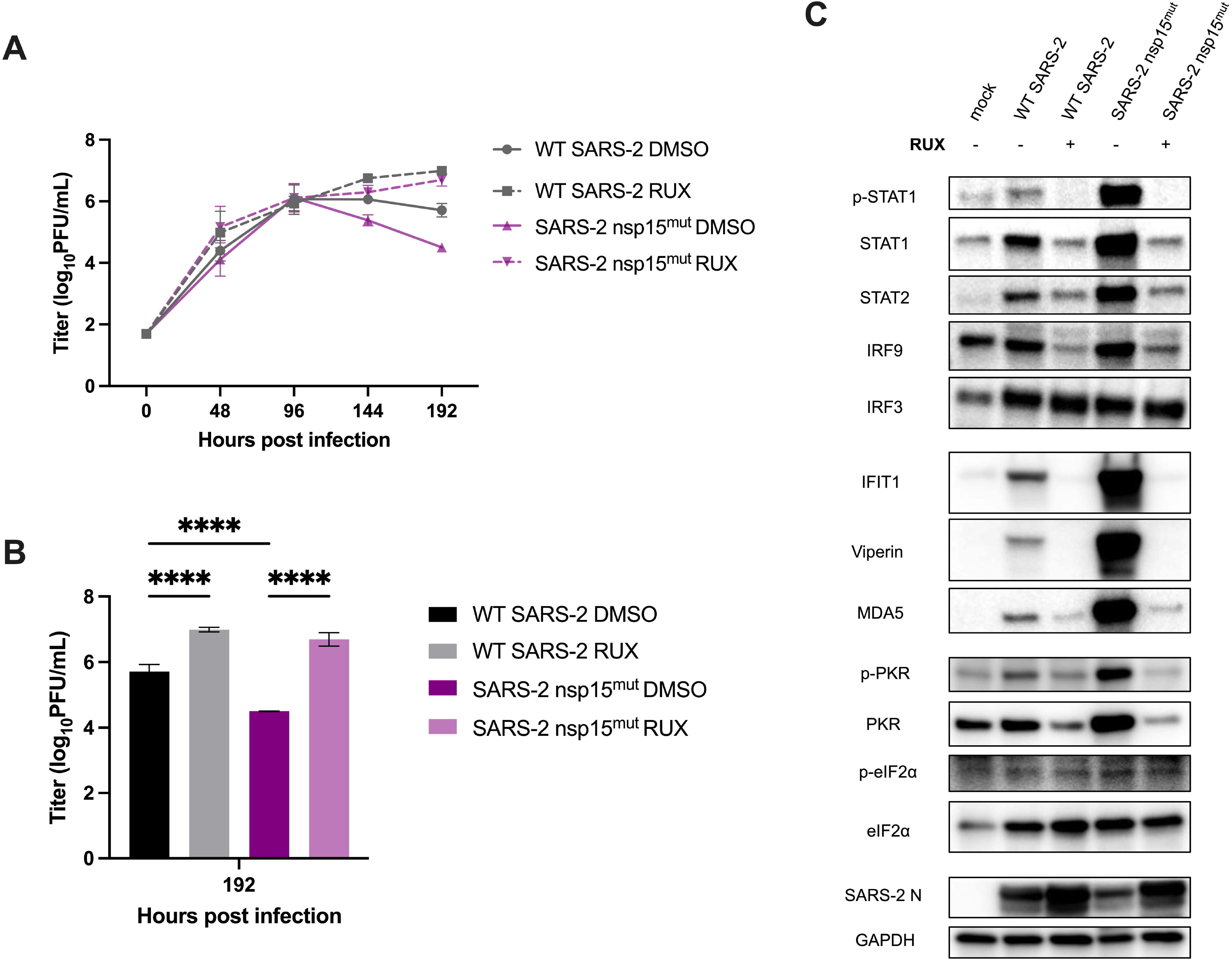
The attenuated replication of SARS-2-nsp15^mut^ relative to WT in primary nasal ALI cultures is IFN-mediated. Primary nasal ALIs were treated with either DMSO or ruxolitinib (RUX) at a concentration of 10 µM for 48 hours prior to infection. Cultures were then infected with either WT or nsp15^mut^ SARS-CoV-2. (A) Apical surface fluid was collected every 48 hours and titered via plaque assay. (B) Represents the 192 hpi data from A presented as a bar graph with significance comparisons shown. (C) Whole cell lysates were collected at 192 hpi for separation via SDS-PAGE and western blot analysis using antibodies against indicated proteins. All data shown is from one experiment representative of two total experiments, and each set of pooled nasal cell cultures is derived from 4-6 individual donors.

Protein lysates from control- or RUX-pretreated infected nasal ALI cultures were analyzed via western blot to confirm the inhibitory activity of RUX and to further investigate the relationships between the PKR pathway and IFN signaling during SARS-CoV-2 infection (**Figure 5C**). At 192 hpi in DMSO-treated control samples, increased ISG expression was observed following SARS-CoV-2 nsp15^mut^ infection compared to WT, as described earlier. RUX treatment completely inhibited STAT1 phosphorylation, as well as upregulation of both STAT1 and STAT2 expression during either WT or nsp15^mut^ SARS-CoV-2 infection. Downstream upregulation of ISG expression was also inhibited in RUX-treated cultures. Interestingly, no increase in p-PKR and total PKR signal was detected in RUX-treated cultures, suggesting that IFN-mediated upregulation of PKR is necessary for PKR pathway activation in nasal ALI cultures. Finally, consistent with viral attenuation, SARS-CoV-2 N levels were decreased during SARS-CoV-2 nsp15^mut^ infection compared to WT in DMSO-treated cultures, but this defect was rescued with RUX treatment. These data indicate that the growth defect observed for SARS-CoV-2 nsp15^mut^ in nasal ALI cultures is IFN-mediated.

### Nsp15 endoribonuclease activity is more crucial in promoting viral replication during MERS-CoV infection compared to SARS-CoV-2 infection

We sought to evaluate the role of nsp15 in innate immune antagonism and optimization of viral replication during SARS-CoV-2 infection compared with another lethal CoV, MERS-CoV. Since these viruses utilize different cellular receptors, we chose two cell types in which both viruses can efficiently replicate for these comparisons (Calu3 and nasal ALI cultures) [50]. For these comparisons we used our previously characterized MERS-CoV mutants, MERS-CoV nsp15^mut^ and a MERS-CoV nsp15^mut^/ΔNS4a double mutant, which also lacks expression of dsRNA-binding protein NS4a. Previously, we found that both of these MERS-CoV mutant viruses were attenuated in A549 cells expressing the MERS-CoV receptor, but the deletion of NS4a in addition to the inactivation of nsp15 conferred a more dramatic replication defect and greater stimulation of IFN signaling and PKR activation [19]. Thus, along with WT SARS-CoV-2 and SARS-CoV-2 nsp15^mut^, we infected Calu3 and nasal ALI cultures with WT MERS-CoV, MERS-CoV-nsp15^mut^ and MERS-CoV-nsp15^mut^/ΔNS4a.

In Calu3 cells, both MERS-CoV nsp15^mut^ and MERS-CoV nsp15^mut^/ΔNS4a were attenuated compared to WT MERS-CoV; this contrasts with the lack of attenuation observed for SARS-CoV-2 nsp15^mut^ (**Figures 6A and 6B**). Additionally, MERS-CoV nsp15^mut^/ΔNS4a exhibited a larger growth defect than MERS-CoV nsp15^mut^ in Calu3 cells. In nasal ALI cultures, WT SARS-CoV-2 replicated to higher titers than WT MERS-CoV (**Figure 7A**), consistent with our previous findings [19, 43]. A significant growth defect for SARS-CoV-2 nsp15^mut^ was observed at 144 and 192 hpi, while both MERS-CoV mutants were attenuated relative to WT MERS-CoV beginning earlier, at 96 hpi. The magnitude of the growth defect was also larger for both MERS-CoV mutants (∼100-fold) compared to SARS-CoV-2 nsp15^mut^ (∼10-fold). Thus, nsp15 activity appears to play a more crucial role in viral replication during MERS-CoV compared to SARS-CoV-2 infection, as MERS-CoV nsp15^mut^ and MERS-CoV nsp15^mut^/ΔNS4a exhibited an earlier and larger replication defect in both cell types.

**Figure 6.**
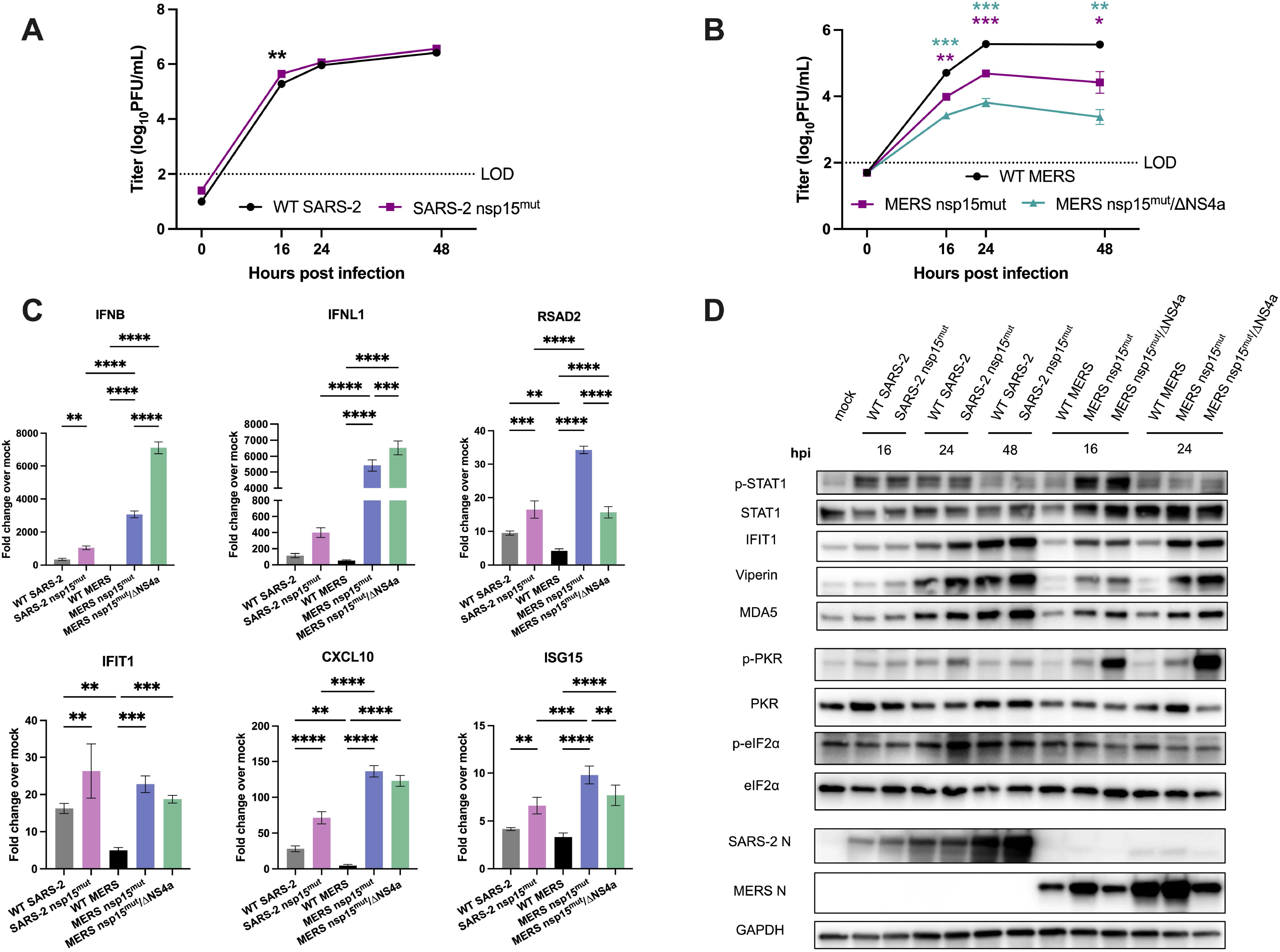
Inactivation of nsp15 endoribonuclease has more impact on viral replication during MERS-CoV infection in Calu3 cells compared to SARS-CoV-2. Calu3 cells were infected with either WT SARS-CoV-2, SARS-CoV-2 nsp15^mut^, WT MERS-CoV, MERS-CoV nsp15^mut^, or MERS-CoV nsp15^mut^/ΔNS4a at an MOI of 0.1. (A-B) Supernatants were collected at the indicated time points post-infection and infectious virus was quantified by plaque assay. (C) Intracellular RNA was extracted at 24 hpi and relative mRNA expression of *IFNB* and *IFNL1* as well as four representative ISGs: *RSAD2*, *IFIT1*, *CXCL10*, and *ISG15* was determined by RT-qPCR. Fold change is expressed as ΔΔCt. (D) Western blot analysis of whole cell lysates was performed at indicated time points using antibodies against indicated proteins. Data shown is from one representative experiment of three total experiments.

**Figure 7.**
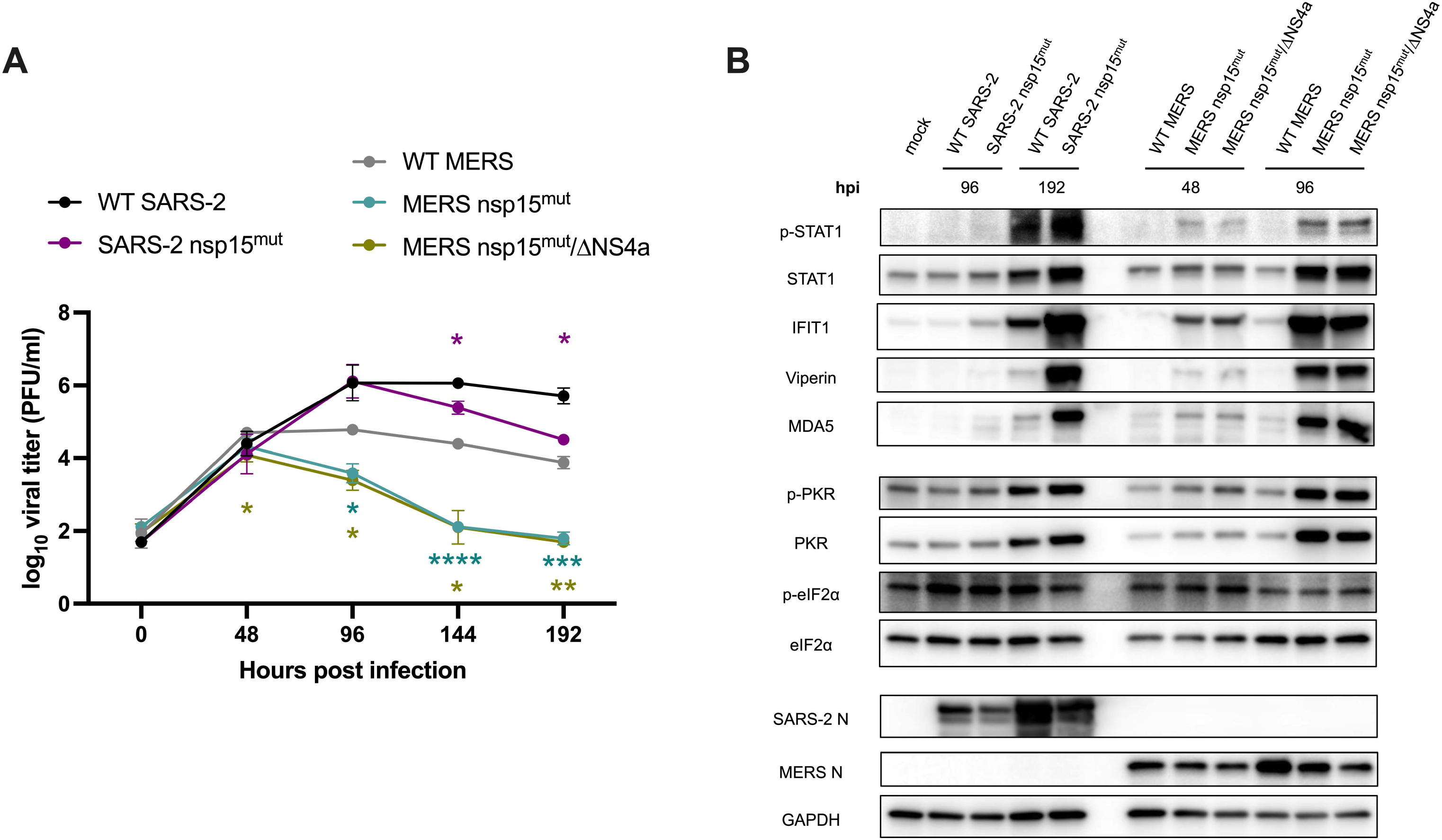
Inactivation of nsp15 endoribonuclease significantly impacts viral replication and IFN signaling of MERS-CoV as well as SARS-CoV-2 in nasal ALI cultures. Nasal ALI cultures were infected with either WT SARS-CoV-2, SARS-CoV-2 nsp15^mut^, WT MERS-CoV, MERS-CoV nsp15^mut^, or MERS-CoV nsp15^mut^/ΔNS4a at an MOI of 1. (A) Apical surface liquid was collected every 48 hours and titered via plaque assay. (B) Whole cell lysates were collected at the indicated time point post-infection for western blot analysis. The blots were probed with antibodies against indicated proteins. All data shown is from one experiment representative of three total experiments, and each set of pooled nasal cell cultures is derived from 4-6 individual donors.

We further compared activation of the IFN and PKR pathways in Calu3 cells. WT MERS-CoV infection resulted in nearly undetectable IFN and ISG mRNA expression whereas WT SARS-CoV-2 infection promoted mild induction of the IFN signaling pathway, consistent with prior experiments (**Figure 6C**) [19, 43]. Infection with SARS-CoV-2 nsp15^mut^ promoted an increase in IFN induction compared to WT SARS-CoV-2 while infection with either MERS-CoV-nsp15^mut^ or MERS-CoV-nsp15^mut^/ΔNS4a induced a more robust increase in *IFNB* and *IFNL1* mRNA expression compared SARS-CoV-2 nsp15^mut^ (>three-fold and >ten-fold, respectively). Interestingly, differences in ISG induction were more nuanced. MERS-CoV mutants induced some ISGs more robustly than SARS-CoV-2 nsp15^mut^ (*CXCL10 and RSAD2*), while other ISGs were induced to similar degrees by SARS-CoV-2 nsp15^mut^ and both MERS-CoV mutants (*IFIT1* and *ISG15*). At the protein level, the SARS-CoV-2 nsp15 mutant as well as both MERS-CoV mutants exhibited peak STAT1 phosphorylation at 16 hpi, although p-STAT1 levels were higher during infection with both MERS-CoV mutants (**Figure 6D**). Both MERS-CoV mutants induced ISGs (IFIT1, viperin, MDA5) earlier (beginning at 16 hpi) than SARS-CoV-2 nsp15^mut^ (first induction at 24 hpi). The SARS-CoV-2 nsp15^mut^ did not reach peak ISG induction levels until 48 hpi. This suggests that nsp15 antagonizes IFN earlier during MERS-CoV infections compared with SARS-CoV-2. Regarding the PKR pathway, neither SARS-CoV-2 nsp15^mut^ nor MERS-CoV-nsp15^mut^ induced PKR phosphorylation as robustly as MERS-CoV-nsp15^mut^/ΔNS4a. This may partially explain why MERS-CoV-nsp15^mut^/ΔNS4a is more attenuated in Calu3 cells.

In nasal ALI cultures, given the delayed IFN induction observed during SARS-CoV-2 infection, we first evaluated IFN and ISG induction for each of the SARS-CoV-2 and MERS-CoV viruses at a late time point (192 hpi). We found that IFN and ISG mRNA expression had returned to mock levels for both MERS-CoV mutants at 192 hpi, suggesting that this time point was too late to detect IFN induction for these viruses (**Figure S4**). Therefore, protein expression of ISGs and PKR pathway activation were examined via western blot at relevant time points for each virus (96, 192 hpi for SARS-CoV-2 and 48, 96 hpi for MERS-CoV) (**Figure 7B**). In line with our observations in Calu3 cells, increased phosphorylation of STAT1 as well as ISG expression occurred at earlier times post-infection in nasal ALI cultures with either of the MERS-CoV mutants compared to SARS-CoV-2 nsp15^mut^. Interestingly, SARS-CoV-2 nsp15^mut^ stimulated the highest accumulation of p-STAT1 at 192 hpi. However, induction of each of the ISGs analyzed occurred to a similar extent with MERS-CoV-nsp15^mut^ and MERS-CoV-nsp15^mut^/ΔNS4a at 96 hpi and SARS-CoV-2 nsp15^mut^ at 192 hpi. Comparable levels of PKR phosphorylation were detected at 48 and 96 hpi with the MERS-CoV mutants and at 192 hpi with SARS-CoV-2 nsp15^mut^. Overall, the kinetics of IFN induction and ISG expression occur at earlier times post infection following inactivation of MERS-CoV nsp15 compared to SARS-CoV-2 nsp15. This kinetic signature correlates with an earlier and more dramatic impact on viral replication during infection with either of the MERS-CoV nsp15 mutants compared to SARS-CoV-2 nsp15^mut^ in nasal ALI cultures. These data indicate that other viral antagonists in addition to nsp15 may be contributing to innate immune antagonism by SARS-CoV-2.

## DISCUSSION

The conserved CoV nsp15 endoribonuclease has been shown to be a potent inhibitor of host innate immunity during infection with multiple CoVs. Infections with viruses expressing nsp15 with an inactive endoribonuclease have resulted in significant attenuation of viral replication compared to their WT counterparts. Increased dsRNA-induced responses, including IFN production and signaling, as well as PKR and OAS/RNase L pathway activation was also observed [5, 16–19]. To build upon previous literature that utilized overexpression systems to identify SARS-CoV-2 nsp15 as an inhibitor of IFN signaling, we generated a recombinant SARS-CoV expressing a catalytically inactive nsp15 EndoU to investigate the role of nsp15 in limiting dsRNA-induced pathway activation and thereby optimizing viral replication.

Despite evidence of robust IFN signatures as well as increased PKR activation in lung-derived epithelial cell lines and primary nasal ALI cultures, replication of SARS-CoV-2 nsp15^mut^ was only attenuated in primary nasal cultures **(Figures 2, 3, and 4)**. These data suggest a role for SARS-CoV-2 nsp15 as an IFN antagonist whose function is essential for optimizing viral replication in certain cellular contexts. Various studies have demonstrated that SARS-CoV-2 is highly sensitive to IFN pre-treatments [51]. Thus, its ability to evade or antagonize IFN and ISG signaling impacts its ability to replicate.

We have previously reported that SARS-CoV-2 infections of nasal ALI cultures produce a relatively low percentage of infected cells (∼10%) compared to cell lines [46]. We hypothesize that this low percentage of infected cells, which may more closely reflect *in vivo* infections, may accentuate the impact that inactivation of nsp15 has on SARS-CoV-2 infection. Secreted IFN from a relatively small population of infected cells provides a signal to neighboring cells to create an antiviral state to limit viral spread [52, 53]. Additionally, nasal ALI cultures are more robust in terms of overall IFN and ISG signaling when compared to transformed epithelial cell lines. We compared the magnitude of IFN and ISG mRNA induction during infection of A549-ACE2, Calu3, and nasal ALI cultures and found that nasal cultures consistently exhibited the strongest induction of *IFIT1* and *ISG15* mRNAs (**Figure S5**). It is also important to note that we observed some variability in the magnitude of IFN and ISG induction within these primary nasal ALI cultures. There is a level of donor-dependent variability in the susceptibility to infection as well as magnitude of immune responses that we and other groups have previously observed (**Figure 4C**) [46, 54]. To help mitigate this variability, four to six individual donors were pooled prior to seeding nasal cultures to model average host responses.

Given that SARS-CoV-2 nsp15^mut^ replication is attenuated in nasal cells, we used a JAK1/2 inhibitor, RUX, to test to what extent the observed growth defect is IFN-mediated. While RUX treatment led to an increase in WT SARS-CoV-2 titers, it had a more robust impact on SARS-CoV-2 nsp15^mut^ titers, such that its replication was rescued to WT SARS-CoV-2 levels in the presence of RUX (**Figure 5**). The correlation between SARS-CoV-2 attenuation and IFN induction is consistent with our previous findings that characterized multiple MERS-CoV nsp15 mutant viruses in A549 cells. We found that growth defects in these mutant viruses were similarly IFN-mediated, as infections conducted in mitochondrial antiviral-signaling protein (MAVS) knockout cells resulted in complete rescue of viral titers to WT MERS-CoV levels in the absence of IFN signaling [19]. Similar findings have been observed with MHV: nsp15 mutant viruses can only replicate efficiently in BMMs deficient in either IFNAR or both PKR and RNase L [17]. Our data further illustrate that nsp15 is an IFN antagonist since SARS-CoV-2 nsp15^mut^ is able to replicate to WT levels when IFN signaling is abrogated in nasal ALI cultures.

RUX treatment of nasal cells also highlighted the interconnected nature of dsRNA-induced pathways. PKR pathway activation indicated by increased p-PKR occurred in nasal cells infected with either WT or nsp15^mut^ SARS-CoV-2. However, RUX treatment resulted in p-PKR levels that were comparable to mock levels during infection by both viruses (**Figure 5**). Since PKR itself is an ISG, this suggested that IFN-mediated upregulation may be necessary for sufficient PKR expression that allows for its activation and autophosphorylation in nasal cell cultures [55, 56]. Parallel observations have been made during infection of BMMs with MHV, whereby low basal expression of OASs resulted in undetectable RNase L activation [57]. In contrast, the downstream target of PKR, eIF2a, is not an ISG and thus would not be upregulated in the context of increased IFN signaling during SARS-CoV-2 nsp15^mut^ infection. Detection of increased phosphorylated eIF2a proved difficult in both Calu3 and nasal cells, which may be linked to its lack of IFN-mediated upregulation. It is important to note that PKR is not the only kinase responsible for eIF2a phosphorylation. Three other kinases (PKR-like ER Kinase (PERK), general control nonderepressible 2 (GCN2), and Heme-regulated eIF2α kinase (HRI)) have the capacity to phosphorylate eIF2a, but only PKR is activated secondary to dsRNA detection. Instead, these kinases are activated by the integrated stress response when an accumulation of unfolded proteins, amino acid starvation, or a heme deficiency is detected [58, 59]. Additionally, we hypothesize that due to the robust activation of both the PKR and RNase L pathways during WT SARS-CoV-2 infection in the epithelial-derived cell lines, any further activation by the nsp15 mutant was difficult to detect.

The IFN-mediated growth defect of the SARS-CoV-2 nsp15^mut^ in nasal ALI cultures was only detected at late times post infection (144 and 192 hpi), concurrent with time points at which the nsp15^mut^ induced more IFN signaling than WT SARS-CoV-2 (**Figure 4**). Similar findings of delayed kinetics of IFN and ISG signaling in primary nasal cells were previously reported based on comparisons with influenza which induces the IFN pathway much earlier than SARS-CoV-2 [60]. The mechanism behind this delay in IFN signaling can likely be explained in part by the immune antagonist activities of accessory and additional nonstructural proteins encoded by SARS-CoV-2. Additional conserved CoV nsps likely contribute to SARS-CoV-2 immune evasion, including nsp1 (which selectively inhibits host protein translation), nsp3 (which modifies host proteins via its macrodomain and deubiquitinase domain to suppress antiviral responses), as well as nsp14 and nsp16 (which contribute to RNA modification important for CoV immune evasion) [61–65]. SARS-CoV-2 encodes multiple additional accessory genes, encoded in ORF3b, ORF6, ORF7a, ORF7b, and ORF8, some of which may play roles in immune evasion [66]. Specifically, the protein encoded by SARS-CoV-2 ORF6 has been reported to inhibit nuclear translocation of transcription factors STAT1/2, thus functioning as an IFN signaling inhibitor [67–70]. Since the SARS-CoV-2 nsp15^mut^ retains other functional IFN antagonist activities in its accessory proteins, this may contribute to the delayed IFN responses observed in the nasal ALI cultures. We previously reported that MERS-CoV encodes three potent IFN antagonists that together shut down dsRNA-induced pathways (the conserved CoV nsp15 EndoU and accessory proteins NS4a and NS4b), so it is likely that SARS-CoV-2 also encodes multiple strategies to evade host innate immunity [19, 21].

Another potential contributor to delayed IFN kinetics observed during SARS-CoV-2 infection and its nsp15 mutant counterpart is temperature. Infections in primary nasal cultures were conducted at 33°C to model *in vivo* nasal airway temperatures, whereas infections in respiratory epithelial cell lines were conducted at 37°C [47, 71]. Temperatures in the nasal passages range from 32-35°C, whereas temperatures in the lung are closer to ambient body temperature (37°C). Temperature has been shown to modulate innate immune responses and replication of SARS-CoV-2 as well as other viruses such as human rhinovirus-16 [48, 72]. Our prior work has indicated that while SARS-CoV-2 can replicate in nasal cultures incubated at either 33°C or 37°C, its replication is optimal at nasal airway temperature (33°C) [46]. This preference for replication of SARS-CoV-2 at 33°C was corroborated in a lower airway ALI model [48]. The delayed kinetics of activation of IFN and ISG signaling in nasal ALI cultures is likely multifactorial, contributed to by both temperature and immune antagonists encoded by SARS-CoV-2.

Having established that nsp15 EndoU activity contributes to IFN antagonism and promotes viral replication during SARS-CoV-2 infection, we directly compared the effects of an inactive nsp15 nsp15 EndoU domain during either SARS-CoV-2 or MERS-CoV infection in Calu3 cells and nasal ALI cultures (**Figures 6 and 7**). MERS-CoV nsp15 mutants exhibited an earlier and more robust IFN signature associated with attenuation in both Calu3 and nasal cultures. While SARS-CoV-2 nsp15^mut^ induced IFN and downstream ISG expression in all cellular systems analyzed, it was only associated with a growth defect in nasal ALI cultures (**Figures 2, 3, and 4**). WT MERS-CoV is particularly adept at shutting down dsRNA-induced pathway activation in contrast to WT SARS-CoV-2, which induces activation of IFN, PKR, and RNase L [43]. Indeed, WT MERS-CoV induced ten-fold less IFN-β and two-fold less IFN-λ mRNA compared to WT SARS-CoV-2 in Calu3 cells (**Figure 6C**). Additionally, although WT SARS-CoV-2 titers were relatively equal to WT MERS-CoV in Calu3 cells, the levels of IFN and ISG induction were all higher during WT SARS-CoV-2 infection (**Figure 6**). We hypothesize that SARS-CoV-2 has evolved to replicate despite moderate innate immune induction, and thus its replication is not significantly impacted when IFN induction is increased above WT levels due to the inactivation nsp15 EndoU. Although nsp15 serves as a strong IFN antagonist encoded by both viruses, our data indicate that nsp15 may play a more crucial role in optimizing MERS-CoV replication. It is not surprising that CoVs would have varying degrees of sensitivity to IFN. SARS-CoV-2, its variants, and SARS-CoV are all genetically similar to each other but demonstrate variability in their sensitivity to IFN [51, 73–76]. Experiments directly comparing the sensitivity of SARS-CoV-2 and MERS-CoV, as well as their respective nsp15 mutants, to IFN would be useful in characterizing these two lethal coronaviruses.

We have characterized a recombinant SARS-CoV-2 nsp15 mutant virus, demonstrating that nsp15 is a potent antagonist of IFN induction and ISG signaling in multiple cellular contexts. Abrogation of IFN antagonism by SARS-CoV-2 nsp15 can have a dramatic impact on viral replication, illustrated most clearly in primary nasal cells in which the SARS-CoV-2 nsp15 mutant exhibits IFN-mediated attenuation of replication. Future studies will investigate the combinatorial role of other SARS-CoV-2 accessory genes that antagonize IFN signaling, such as the ORF6 encoded protein, as well as the downstream consequences of increased dsRNA-induced pathway activation, such as inflammatory cytokine production and cell death, and how these responses may contribute to overall pathogenesis. Additionally, although nasal ALI cultures possess a heterogenous cellular population of ciliated and mucus-secreting epithelial cells, they lack immune cells, which have been shown to play a prominent role in pathogen recognition as well as inflammatory cytokine production [77]. The establishment of a co-culture nasal ALI system containing various innate immune cell populations would mitigate this pitfall present in all three cellular systems used in this study. It is imperative to characterize the mechanisms of innate immune evasion by SARS-CoV-2 and other HCoVs to understand how viruses interact with host cells to optimize replication and spread. Moreover, these findings will inform discovery of effective antivirals and development of live-attenuated vaccines against SARS-CoV-2 and other respiratory coronaviruses.

## MATERIALS & METHODS

Nasal epithelial cells were collected via cytologic brushing of patients’ nasal cavities after obtaining informed consent and then grown and differentiated on transwell inserts to establish air-liquid interface (ALI) cultures). The full study protocol was approved by the University of Pennsylvania Institutional Review Board (protocol # 800614) and the Philadelphia VA Institutional Review Board (protocol #00781). A549-ACE2, Calu3, and primary nasal epithelial cells were infected with recombinant WT SARS-CoV-2, SARS-CoV-2 nsp15^mut^, WT MERS-CoV, MERS-CoV nsp15^mut^, or MERS-CoV nsp15^mut^/ΔNS4a at the indicated multiplicities of infection [19, 43]. At various times post infection, infectious virus in cellular supernatants or in apical surface liquid collected via apical wash of nasal ALI cultures were quantified via plaque assay. Total RNA or protein was also collected from infected cells for analysis of dsRNA-induced pathway activation. All of these techniques are described in SI Appendix, Materials & Methods. Any materials or related protocols mentioned in this work can be obtained by contacting the corresponding author.

## Acknowledgments

We thank members of the Weiss lab for feedback and discussion of this project and Dr. Volker Thiel (University of Bern, Switzerland) for discussion and early mutant construction. We thank Drs. David W. Kennedy, James N. Palmer, Nithin D. Adappa, and Michael A. Kohanski for aid in the collection of nasal tissue for establishing primary nasal epithelial cultures. This work was supported by National Institutes of Health grants R01 AI140442 (SRW), R01AI169537 (SRW&NAC), R01AI104887 (SRW&RHS), R35GM138029 (ARF), P20GM113117 (ARF); RO1A1AI161175 (LM-S); Department of Veterans Affairs Merit Review 1-I01-BX005432-01 (NAC&SRW); the Penn Center for Research on Coronaviruses and Other Emerging Pathogens (SRW). CO was supported in part by F30AI172101 and T32AI055400 and NB in part by T32AI007324 and K12GM081259.

## DISCLOSURES

Susan R Weiss is on the Scientific Advisory Board of Ocugen, Inc. and consults for Powell Gilbert LLP. Noam A Cohen consults for GSK, AstraZeneca, Novartis, Sanofi/Regeneron; has US Patent “Therapy and Diagnostics for Respiratory Infection” (10,881,698 B2, WO20913112865) and a licensing agreement with GeneOne Life Sciences.

