## Supplementary material for "SARS-CoV-2 nsp15 endoribonuclease antagonizes dsRNA-induced antiviral signaling": SI Appendix, Materials and Methods

### MATERIALS & METHODS

#### Recombinant viruses

Recombinant SARS-CoV-2 viruses were derived from a bacterial artificial chromosome (BAC) vector containing the full-length SARS-CoV-2 USA-WA1/2020 genome. For SARS-COV-2 nsp15$^{mut}$ nucleotide 20,320 (C) and 20,321 (A) were mutated in order to substitute histidine 234 with alanine, and mutant was rescued as previously described [1, 2]. MERS-CoV (HCoV-EMC/2012) was also derived from a BAC. MERS-CoV-nsp15$^{H231A}$ and MERS-CoV-nsp15$^{H231A}$/ΔNS4a were generated as previously described [3]. Virus stocks were sequenced and compared to the publicly available wild-type sequences on NCBI. All virus stocks were generated via low MOI infections in VeroE6 cells (for SARS-CoV-2 viruses) and VeroCCL81 cells (for MERS-CoV viruses).

#### Cloning and protein purification

SARS-CoV-2 nsp15 wild type and H234A mutant was PCR amplified using cDNA clones with forward primer 5'-TTCAAGGATCCATGAGTTTAGAAAATGTGGCTTTTAATG-3' (with BamHI restriction site [underlined]) and reverse primer 5'-TTCAAGTCGACCTATTGTAATTTTGGGTAAAATGTTTC-3' (with SalI restriction site [underlined]). The SARS-CoV-2 nsp15 wild type cDNA template was a gift from Prof. Wang Pei-Hui (Shandong University, China). H234A mutant cDNA clone was obtained by lambda red recombination of the full-length BAC and restriction cloning into the expression vector [4]. The amplified DNA product was cloned into pGEX-6P1 vector at BamHI and SalI site. Nsp15 purification was performed by modifying methods previously described for glutathione S-transferase (GST) fusion proteins [5, 6]. Briefly, GST fusion proteins were expressed from pGEX-6P-1 constructs in *E. coli* strain BL21(DE3)/pLysS (ThermoFisher). Secondary cultures were grown to an optical density (OD) (at 600 nm) of 0.6 in a shaking incubator at 37°C at 250 rpm.

Cells were induced with 0.2 mM isopropyl-β-d-thiogalactopyranoside (IPTG) for 16 hours at 22°C. Induced cell pellets were harvested and suspended in buffer A (20 mM HEPES (pH 7.5), 1 M KCl, 1 mM EDTA, 5 mM dithiothreitol (DTT), 10% (v/v) glycerol, and EDTA-free Pierce protease inhibitor (ThermoFisher)). Pelleted cells were lysed by the addition of 200 µg/ml lysozyme, and sonication. Supernatants were added to Pierce glutathione agarose (ThermoFisher) and incubated for 2 hours at 4°C, followed by washes with buffer A. Digestions to cleave the GST tag were performed with PreScission protease (Cytiva) in a solution containing 50 mM Tris-HCl (pH 7.5), 150 mM NaCl, 1 mM EDTA, and 1 mM DTT for 16 hours at 4°C. Concentrated supernatants containing tag-less protein were loaded onto a Superdex 75 column on an Äkta pure 25L protein purification system (GE Healthcare) in a buffer containing 20 mM HEPES (pH 7.5), 150 mM NaCl, and 1 mM DTT. Protein concentrations were estimated using the Bio-Rad protein assay reagent (Bio-Rad). All proteins were stored in buffer supplemented with 10% glycerol at −80°C until further use.

**Endonuclease activity assay**

Endonuclease activity of nsp15 was determined in a fluorescence based kinetic assay. Synthetic RNA substrate (6-FAM-UUA UCA AAU UCU UAU UUG CCC CAU UUU UUU GGU UUA-BHQ-1) with a FRET (Fluorescence resonance energy transfer) pair (6-FAM at the 5′-terminus and a black hole quencher-1 (BHQ1) at the 3′-terminus) was commercially obtained from IDT [7]. Final reaction mixtures contained 20 mM HEPES (pH 7.5), 1 mM DTT, 5 mM $MnCl_2$, 0.2 µM RNA substrate in the absence or presence of 50 nM wild type or  nsp15 in a 50 µl volume. The reactions were set in a round bottom black polystyrene 96 well plate (Costar) and briefly spun. The reaction kinetics was monitored by measuring fluorescence using a Varioskan LUX microplate reader (ThermoFisher) set at an excitation/emission wavelengths of 485nm/535nm (excitation bandwidth

12 nm and measurement time 100 ms). The kinetics were recorded at intervals of 5 min over a period of 60 min using SkanIt Software 6.0.1.

**Cell lines**

Vero E6 cells and CCL81 Vero cells (ATCC) were cultured in Dulbecco's modified Eagle's medium (DMEM) containing 10% L-glutamine and 4.5g/L D-glucose (Gibco, ThermoFisher) was supplemented with 10% heat inactivated (HI) fetal bovine serum (FBS) (Hyclone, Cytiva), 1X penicillin/streptomycin (pen/strep) (Gibco, ThermoFisher). Human A549 cells engineered to stably express angiotensin-converting enzyme 2 (ACE2) (A549-ACE2) were cultured in RPMI 1640 (Gibco, ThermoFisher) supplemented with 10% HI FBS, and 1X pen/strep [8]. Human Calu3 cells were (ATCC) cultured in DMEM containing 10% L-glutamine and 4.5g/L D-glucose supplemented with 20% HI FBS, 1X pen/strep.

**Nasal air-liquid interface (ALI) culture growth and differentiation**

Nasal specimens were obtained via cytologic brushing of patients in the Department of Otorhinolaryngology-Head and Neck Surgery, Division of Rhinology at the University of Pennsylvania and the Philadelphia Veteran Affairs Medical Center after obtaining informed consent. Patients with any history of systemic disease and those currently on immunosuppressive conditions were excluded. The full study protocol, including acquisition and use of nasal specimens, was approved by the University of Pennsylvania Institutional Review Board (Protocol #800614) and the Philadelphia VA Institutional Review Board (Protocol #00781). ALI cultures were grown on semi-permeable transwell supports (STEMCELL Technologies) (0.4 $\mu$m pores) as previously described [9-11]. Nasal cells derived from 4-6 patients were pooled prior to seeding on to transwell inserts (STEMCELL Technologies). Nasal cells were grown to confluence with

Pneumacult-Ex Plus growth medium (STEMCELL Technologies) present both apically and basally until reaching confluence. At that point apical growth medium was removed. Pneumacult-ALI medium (STEMCELL Technologies) was used to differentiate all nasal ALI cultures and was replaced two times per week for 4 weeks prior to infection. Growth and differentiation of cultures was conducted at 37ºC.

**Infections in A549-ACE2 and Calu3 cell lines**

Viruses were diluted in serum-free (SF) DMEM or RPMI and added to cells for absorption for 1 hour at 37°C. All infections were performed at an MOI of 1 unless otherwise indicated. Cells were washed three times with phosphate-buffered saline (PBS) after the incubation period and DMEM supplemented with 4% HI FBS for Calu3 infections or RPMI supplemented with HI 2% FBS for $A549^{ACE2}$ infections was added. 200μL of supernatant were collected at the indicated time points for infectious virus quantification. The third PBS wash was collected and represents the zero hour time-point to quantify remaining input virus. All infections, plaque assays, and virus manipulations were carried out in a biosafety level 3 laboratory utilizing appropriate personal protective equipment and protocols.

**Infections in nasal ALI cultures**

After differentiation, nasal ALI cultures were allowed to equilibrate at 33ºC for at least 24 hours prior to infection in order to replicate conditions of the *in vivo* nasal airway as previously described [11, 12]. In brief: Viruses were diluted in SF DMEM to achieve a total inoculum volume of 60 μL at an MOI of 1. The inocula were added apically to nasal cultures for a 1 hour adsorption period. After 1 hour, cells were washed on the apical surface three times with PBS. The third PBS wash was collected and represents the 0-hour time point. Every 48-hours following infection, 200 μL PBS was added apically to each nasal ALI culture in order to collect shed virus for quantification.

**Infectious virus quantification**

Extracellular supernatants were titer via plaque assay on either Vero E6 cells for SARS-CoV-2 or Vero CCL81 cells for MERS-CoV using previously described plaque assay methods (Fausto et al., 2023). Briefly, the indicated cell type was seeded ~24 hours before use. The next day, 1:10 serial dilutions of viral supernatants were performed in SF DMEM. Growth media was removed from each well and selected dilutions were added. Plates were incubated for 1 h at 37 ºC. At the end of the 1 hour incubation, enough liquid overlay (DMEM with L-Glut (ThermoFisher) supplemented with 2% HI FBS, 1% sodium pyruvate (ThermoFisher) and 0.01% molecular grade agarose (Thermofisher)) to thoroughly cover each well was added. Plates were incubated at 37 ºC for 3 days for both SARS-CoV-2 and MERS-CoV. After 3 days, the overlay was removed and 4% paraformaldehyde (PFA) in PBS (ThermoFisher) was added to each well for at least 30 min to fix the monolayer and inactivate the virus. The PFA was then removed and disposed of appropriately. The plates were stained using crystal violet (1% crystal violet (Sigma) in 20% ethanol (Fisher Scientific) and 80% water) and rinsed with water before counting. Each sample was plated in technical duplicate. Titer (plaque forming unit (PFU) per ml) represents the average plaque count divided by the dilution factor and inoculum volume in ml.

**Reverse transcriptase (RT)-quantitative PCR**

At indicated times post-infection cells were lysed with buffer RLT Plus (Qiagen RNeasy Plus #74136) and RNA extracted following the manufacturer's protocol. Methods were adapted from Fausto et al. [13]. RNA was then reverse transcribed to make cDNA using a High-Capacity cDNA Reverse Transcriptase Kit (Applied Biosystems, ThermoFisher) (200μg RNA per 20uL cDNA reaction). cDNA was amplified using specific PCR primers (see Table S2), iQ™ SYBR® Green

Supermix (Bio-Rad), and the QuantStudio™ 3 PCR system (Thermo Fisher). Fold changes in ISG mRNA level relative to mock-infected samples were calculated using the formula $2^{-\Delta(\Delta Ct)}$ ($\Delta Ct = Ct_{\text{gene of interest}} - Ct_{18S}$) and expressed as fold changes over mock. Primer sequences for each gene analyzed are provided in Table S2. SARS-CoV-2 genome copy numbers were quantified with primers directed against nsp12 (RdRp) and calculated using a standard curve generated with a digested plasmid encoding SARS-CoV-2 nsp15.

**Western blot analysis**

Cell lysates were harvested at indicated times post infection with lysis buffer (50 mM Tris hydrochloride pH 8.0, 150mM sodium chloride, 0.5% deoxycholate, 0.1% sodium dodecyl sulfate (SDS), 1% nonyl phenoxypolyethoxylethanol (NP)-40, 0.02% sodium azide) supplemented with protease inhibitor cocktail (cOmplete mini EDTA-free protease inhibitor tablets, Roche) and phosphatase inhibitors (PhosSTOP phosphatase inhibitor cocktail tablets, Roche). Lysates were mixed 3:1 with 4x Laemmli sample buffer (Bio-Rad). Samples were heated to 95°C for 10 minutes, resolved on gradient SDS/PAGE gels (Bio-Rad), and subsequently transferred to a polyvinylidene difluoride (PVDF) membrane (Bio-Rad). Blots were blocked with 5% non-fat milk or 5% bovine serum albumin (BSA) in Tris buffered saline with 0.1% Tween 20 (TBST) and probed with the antibodies at the specified concentrations in their respective blocking buffers listed in **Table 2**. Blots were visualized using SuperSignal West Femto Chemiluminescent substrate (ThermoFisher). Blots were stripped using Restore Western Blot Stripping Buffer (ThermoFisher) for at least 1 hour at room temperature. After stripping, blots were thoroughly washed with TBST and blocked prior to the addition of the next primary antibody.

**Ruxolitinib treatments**

48 hours prior to infection, basal media of nasal ALI cultures was supplemented with 10 μM ruxolitinib (Selleck Chem) resuspended in DMSO. Control cultures were treated with DMSO. Basal media containing fresh ruxolitinib was replaced at 0, 48, and 96 hpi. Sample collection and analysis was performed as described here.

**Analyses of RNase L-Mediated rRNA degradation**

Intracellular RNA was harvested with RLT plus buffer and extracted using Qiagen RNeasy Plus kit. RNA was prepped and analyzed on a RNA chip with the Agilent Bioanalyzer using the Agilent RNA 6000 Nano Kit according to manufactures recommendations [14].

**Statistics**

Data was graphed and statistics were performed using GraphPad Prism. Data was graphed displaying either individual values or the mean +/- the standard deviation. Statistical significance was determined by comparing mutant viruses to WT using either a one-way ANOVA for single time point comparisons and a two-way ANOVA for experiments with multiple time points and multiple viruses. Significance shown represents the P values where * = $P < 0.05$; ** = $P < 0.01$; *** = $P < 0.001$; and **** = $P < 0.0001$; ns = not significant. In some figures, ns is not displayed on the graph.

**Table S2. Primers used for qPCR analysis**

| Gene name | Primer orientation | Sequence |
|---|---|---|
| IFNL1 | Forward | CGCCTTGGAAGAGTCACTCA |
| | Reverse | GAAGCCTCAGGTCCCAATTC |
| IFNB | Forward | GTCAGAGTGGAAATCCTAAG |
| | Reverse | ACAGCATCTGCTGGTTGAAG |
| IFIT1 | Forward | TGGTGACCTGGGGCAACTTT |

| | | |
|---|---|---|
| | Reverse | AGGCCTTGGCCCGTTCATAA |
| RSAD2 | Forward | CACAAAGAAGTGTCCTGCTTGGT |
| | Reverse | AAGCGCATATATTCATCCAGAATAAG |
| CXCL10 | Forward | CCTGCAAGCCAATTTTGTCC |
| | Reverse | ATGGCCTTCGATTCTGGATTC |
| ISG15 | Forward | CATCTTTGCCAGTACAGGAGC |
| | Reverse | GGGACACCTGGAATTCGTTG |
| 18S | Forward | TTCGATGGTAGTCGCTGTGC |
| | Reverse | CTGCTGCCTTCCTTGAATGTGGTA |
| SARS-CoV-2 nsp12 (RdRp) | Forward | GGTAACTGGTATGATTTCG |
| | Reverse | CTGGTCAAGGTTAATATAGG |

**Table S2. Antibodies used for western blotting**

| Primary Antibody | Antibody species | Blocking buffer | Dilution | Catalog number |
|---|---|---|---|---|
| p-PKR | Rabbit | 5% BSA in TBST | 1:1000 | Abcam 32036 |
| PKR | Rabbit | 5% milk or 5% BSA in TBST | 1:1000 | Cell Signaling 12297 |
| p-eIF2α | Rabbit | 5% BSA in TBST | 1:1000 | Cell Signaling 3398 |
| eIF2α | Rabbit | 5% milk or 5% BSA in TBST | 1:1000 | Cell Signaling 9722 |
| IFIT1 | Rabbit | 5% milk or 5% BSA in TBST | 1:1000 | Cell Signaling 14769 |
| Viperin | Rabbit | 5% milk or 5% BSA in TBST | 1:1000 | Cell Signaling 13996 |
| MDA5 | Rabbit | 5% milk or 5% BSA in TBST | 1:1000 | Cell Signaling 5321 |
| p-STAT-1 | Rabbit | 5% BSA in TBST | 1:1000 | Cell Signaling 7649 |
| STAT1 | Rabbit | 5% milk or 5% BSA in TBST | 1:1000 | Cell Signaling 9172 |

| | | | | |
|---|---|---|---|---|
| STAT2 | Rabbit | 5% milk or 5% BSA in TBST | 1:1000 | Cell Signaling 72604 |
| IRF9 | Rabbit | 5% milk or 5% BSA in TBST | 1:500 | Santa Cruz SC-10793 |
| IRF3 | Rabbit | 5% milk or 5% BSA in TBST | 1:1000 | Cell Signaling 11904 |
| SARS-CoV-2 Nucleocapsid | Rabbit | 5% milk or 5% BSA in TBST | 1:2000 | Genetex GTX135357 |
| MERS-CoV Nucleocapsid | Mouse | 5% milk or 5% BSA in TBST | 1:2000 | Sino Biological 40068-MM10 |
| GAPDH | Rabbit | 5% milk or 5% BSA in TBST | 1:2000 | Cell Signaling 2118 |
| **Secondary Antibody** | **Antibody species** | **Blocking buffer** | **Dilution** | **Catalog number** |
| Anti-rabbit IgG HRP-linked | Goat | Same as primary | 1:3000 | Cell Signaling 7074 |
| Anti-mouse IgG HRP-linked | Horse | Same as primary | 1:3000 | Cell Signaling 7076 |

**Figure Legends for Supplement**

**Figure S1. Purification of WT and H234A mutant nsp15 proteins**

Each recombinant protein was purified and visualized on a gel to confirm homogeneity. See cloning and protein purification section of Materials & Methods for further detail.

**Figure S2. SARS-CoV-2 nsp15$^{mut}$ does not induce the OAS/RNase L pathway above WT SARS-CoV-2 levels in A549-ACE2 cells**

A549-ACE2 cells were infected at an MOI of 1 with either WT or nsp15$^{mut}$ SARS-CoV-2. Total cellular RNA was collected at the indicated time points post infection and were analyzed for RNase L activation. Data shown are from one representative experiment of three experiments.

**Figure S3. SARS-CoV-2 nsp15$^{mut}$ does not induce the OAS/RNase L pathway above WT SARS-CoV-2 levels in Calu3 cells**

Calu3 cells were infected at MOI of 1 with WT or nsp15$^{mut}$ SARS-CoV-2 . Total cellular RNA was collected at indicated times post infection and were assessed for RNase L activity.

**Figure S4. IFN and ISG mRNA expression by SARS-CoV-2 and MERS-CoV mutants at 192 hours post-infection in nasal ALI cultures**

Nasal cell cultures were infected at MOI of 1 with either WT SARS-CoV-2, SARS-CoV-2 nsp15$^{mut}$, WT MERS-CoV, MERS-CoV nsp15$^{mut}$, or MERS-CoV nsp15$^{mut}$/∆NS4a in triplicate. Total cellular RNA was collected from infected cultures at 192 hpi and mRNA expression analysis for *IFNL1, IFNB, IFIT, CXCL10, RSAD2,* and *ISG15* using the ∆∆Ct formula to calculate fold changes in expression relative to mock cultures, normalized to *18S*.

**Figure S5. Comparison of fold change induction of IFN and ISG mRNAs in A549-ACE2 cells, Calu3 cells, and nasal ALI cultures**

Cells from the indicated culture system were infected with either WT or nsp15$^{mut}$ SARS-CoV-2. RNA was collected in triplicate for analysis of fold change in mRNA expression above mock-infected levels, calculated using $\Delta\Delta$Ct formula for indicated IFN or representative ISG mRNA. Fold changes from 3 independent experiments (triplicate values from each experiment) were plotted for the time point of maximal fold change induction in IFN and ISG expression in that cellular system (72 hpi for A549-ACE2, 24 hpi for Calu3, 192 hpi for nasal ALI cultures). Fold changes over mock are plotted on the same graph for each gene in order to compare degree of induction in each cellular system.

**S1**

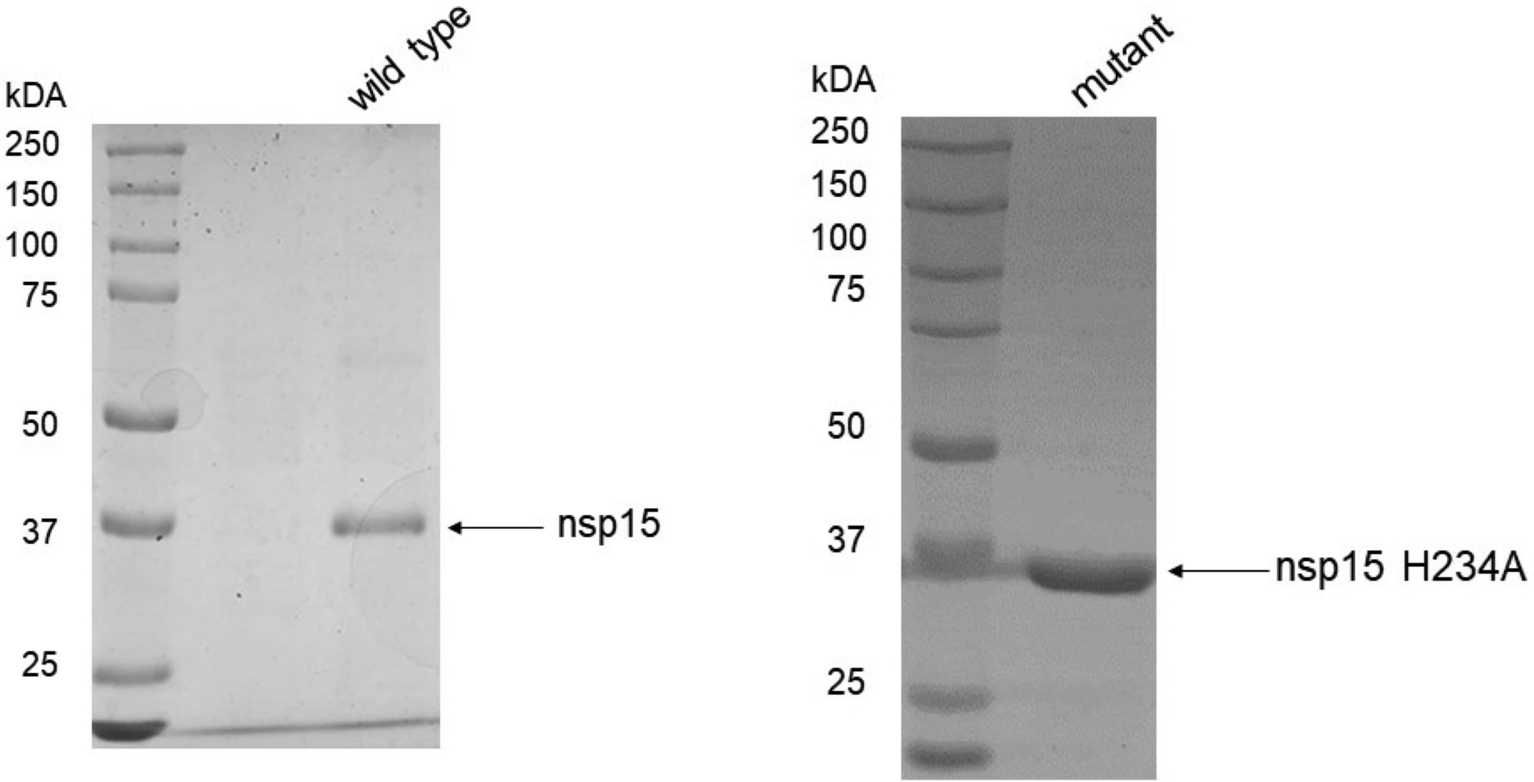

wild type
kDA
250
150
100
75
50
37
25
nsp15
mutant
kDA
250
150
100
75
50
37
25
nsp15 H234A

**S2**

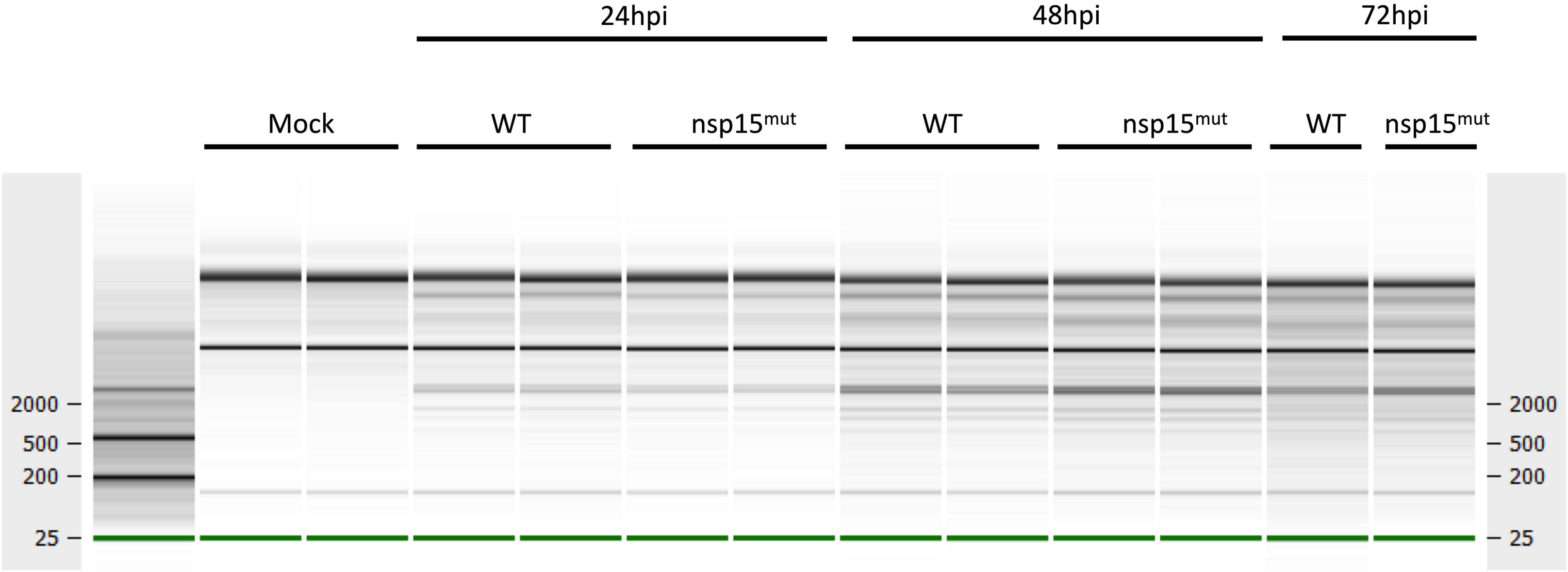

24hpi
48hpi
72hpi
Mock
WT
nsp15mut
WT
nsp15mut
WT
nsp15mut
2000
500
200
25
2000
500
200
25

S3

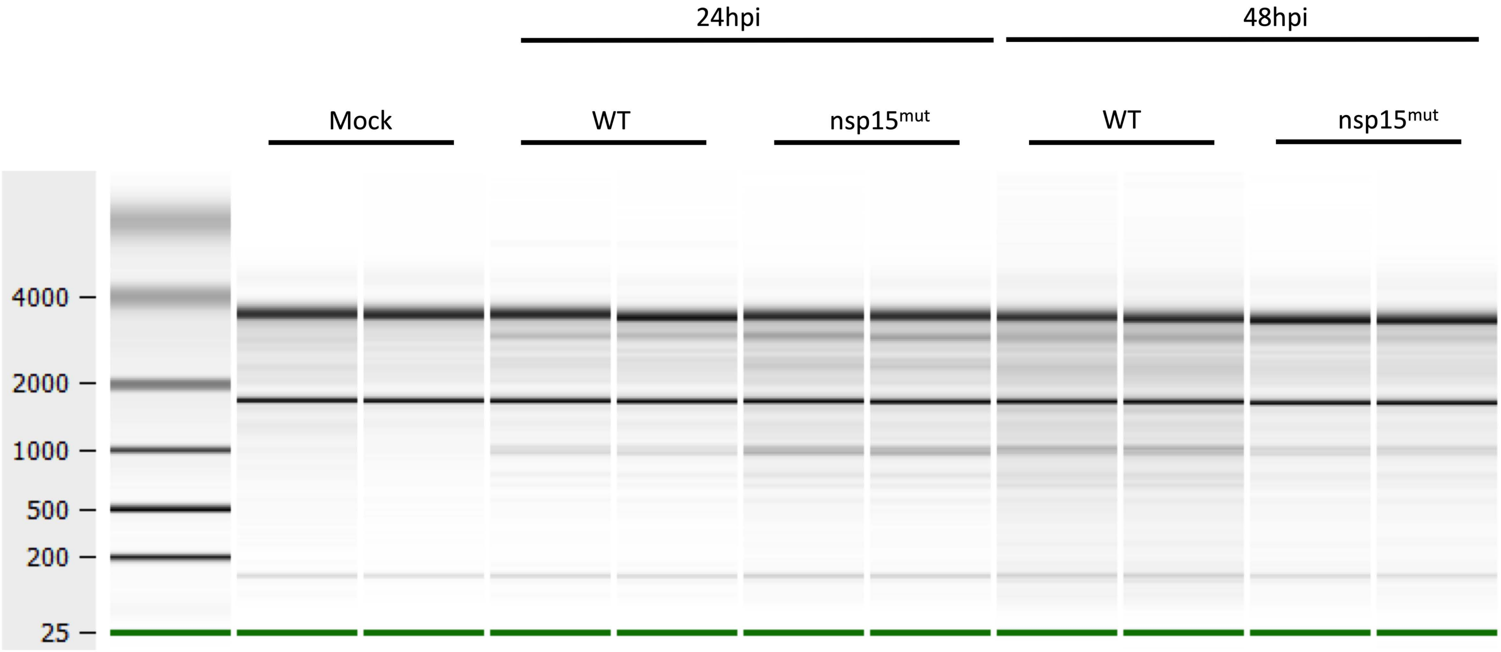
24hpi
48hpi
Mock
WT
nsp15mut
WT
nsp15mut
4000
2000
1000
500
200
25

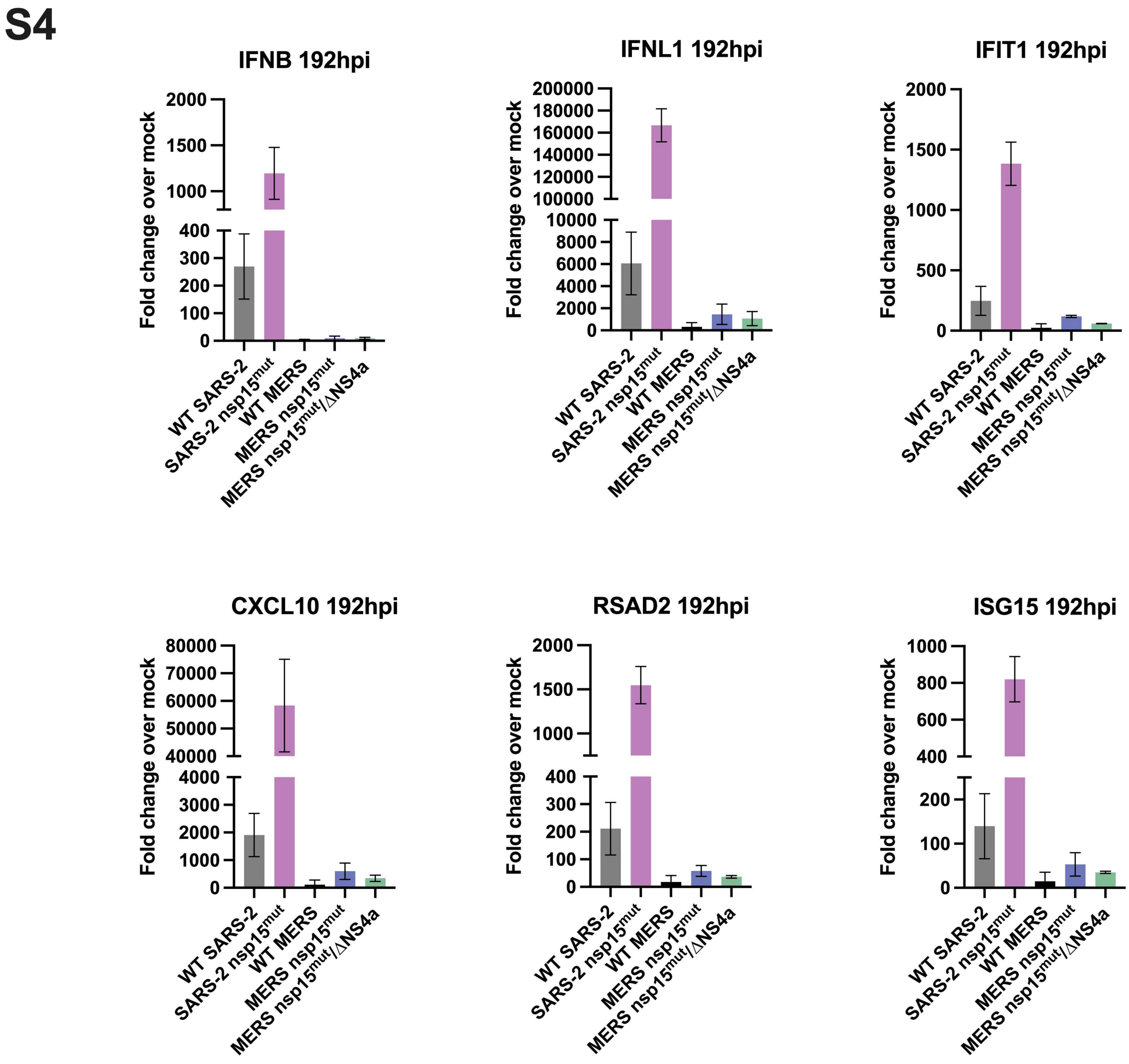

S4
IFNB 192hpi
IFNL1 192hpi
IFIT1 192hpi
CXCL10 192hpi
RSAD2 192hpi
ISG15 192hpi
Fold change over mock
WT SARS-2
SARS-2 nsp15mut
WT MERS
MERS nsp15mut
MERS nsp15mut/ΔNS4a

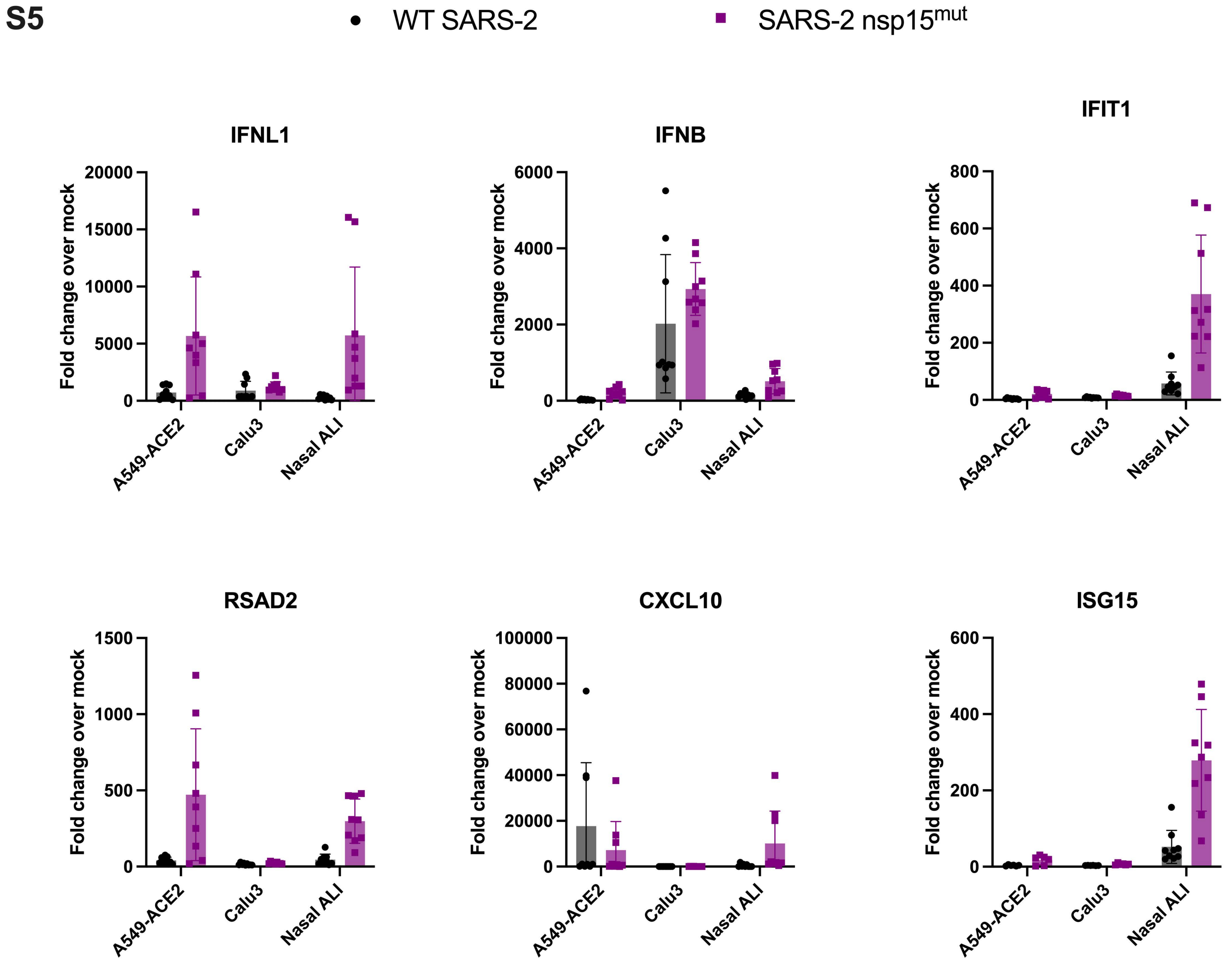
S5
WT SARS-2
SARS-2 nsp15mut
IFNL1
IFNB
IFIT1
RSAD2
CXCL10
ISG15
Fold change over mock
A549-ACE2
Calu3
Nasal ALI
20000
15000
10000
5000
0
6000
4000
2000
800
600
400
200
1500
1000
500
100000
80000
60000
40000
20000
